# Adipocyte Expression of O-Glycoprotein Procollagen C-Endopeptidase Enhancer Protein 2 (PCPE2): Mechanisms Linking Fibrosis and Beiging of White Adipose Tissue

**DOI:** 10.64898/2026.07.24.740318

**Authors:** Mirza A Beg, Hao Xu, Bilal Ahmad, Alexander Rocksvold, Rana K Gupta, Pablo Esteban Morales, Ze Zheng, Wen Dai, Justin Grobe, John J. Reho, Yiliang Chen, Samuel Klein, Gordon I. Smith, Jong Kyoung Kim, Eun Seo Park, Hahn Nahmgoong, Jae Bum Kim, Deniz Ince, Stacy A. Malaker, Rachel L. Mintz, Gwendalyn J Randolph, Mary G. Sorci-Thomas

## Abstract

**Background:** Procollagen C-endopeptidase enhancer protein 2 (Pcpe2) has been primarily investigated in collagen processing during wound healing and is assumed to function similarly to Pcpe1 in the extracellular matrix (ECM). Our studies suggest that Pcpe2 has unique structural and functional features not shared with Pcpe1.

**Methods:** To study the role of Pcpe2 in adipose tissue remodeling, we created adipose tissue-specific knockout of Pcpe2 (TgAd+Pcpe2KO) and adipose tissue-specific Pcpe2 overexpressing (TgAd+Pcpe2Ox) mice and performed in vivo and ex vivo experiments.

**Results:** We show that TgAd+Pcpe2KO mice are resistant to WD-induced obesity and exhibit reductions in body and fat pad mass. Promethion cage studies showed no significant differences in lean mass or food intake, yet WD-fed TgAd+Pcpe2KO mice exhibited significantly higher energy expenditure than negative Cre littermate controls. WD-fed TgAd+Pcpe2KO mice also showed reduced plasma glucose and lipoprotein lipid concentrations compared to controls. While markers of adipose tissue inflammation were reduced in WD-fed TgAd+Pcpe2KO mice. Examination of mature adipocytes from white adipose tissue showed that WD consumption greatly stimulated Pcpe2 expression, while no change in Pcpe2 expression in the stromal vascular cells was noted. However, in VAT precursor cells (PCs), the CD140b+ population showed a shift from fibroadipogenic precursors towards adipocyte PCs in TgAd+Pcpe2KO mice. This shift in PCs may account for the attenuation of local inflammation and fibrosis in the absence of Pcpe2. Ex vivo differentiation of adipose PCs showed that loss of Pcpe2 enhanced adipocyte differentiation, reducing TGFβ-like signaling via pSmad2/3, and increasing mitochondrial function. Furthermore, unlike Pcpe1, Pcpe2 contains a mucin-like linker domain with nine sites of O-linked glycosylation which may regulate receptor signaling at the plasma membrane.

**Conclusions:** Our results show the ECM O-glycoprotein Pcpe2 is a robust marker of unhealthy adipose tissue expansion in humans and mice and contributes to inflammation and fibrosis associated with WD-induced obesity.

## Introduction

Obesity is a chronic disease contributing to life-limiting metabolic comorbidities, including type 2 diabetes,^1^ cardiovascular disease,^2^ and steatotic liver disease.^3^ Excessive caloric intake can lead to obesity, which is characterized by dysfunctional lipid storage and chronic inflammation.^4^ Pathogenesis associated with obesity is closely linked to the inability of white adipose tissue (WAT) to efficiently store lipid in adipocytes.^5,6^ Adipose tissue acts as a central metabolic hub, storing fat for energy and mediating systemic insulin sensitivity.^7,8^ However, the constant demand to expand WAT leads to adipocyte hypertrophy and reduced production of newly differentiated adipocytes.^9^ Excessive fat storage within existing adipocytes leads to chronic, low-grade inflammation and ectopic fat accumulation, thereby driving insulin resistance and systemic metabolic decline.^10–12^ Therefore, identifying novel molecular regulators and understanding the mechanisms governing adipose tissue health, particularly those involved in inflammation, adipocyte differentiation, and energy expenditure, is paramount for developing effective therapeutic strategies against obesity.

WAT remodeling is a highly dynamic process essential for metabolic adaptation, allowing adipocytes to expand or contract in response to nutritional changes, such as energy surplus or deficit.^8,13^ This process involves coordinated cellular dynamics, including changes in adipocyte size and number, and requires the active participation of various cells in the stromal vascular fraction (SVF) in WAT.^14,15^ In adipose tissue, there are mature adipocytes as well as precursor cells (PCs) in the SVF,^16,17^ which, given appropriate signals, differentiate to mature adipocytes.^15,18^ Adipocyte precursor cells (APCs) and fibroadipogenic progenitors (FAPs)^18,19^ are located in visceral adipose tissue (VAT) SVF and are uniquely responsive to diet.^20,21^ Under metabolic stress, after consuming a Western diet (WD), FAPs secrete cytokines that inhibit the differentiation of APCs into mature adipocytes.^22,23^ These pro-inflammatory signals stimulate continued macrophage infiltration and exacerbate inflammation,^24^ leading to elevated expression of interleukin-6 (Il-6), C-C motif chemokine ligand 2 (Ccl2), tumor necrosis factor alpha (Tnf-α), and hypoxia-inducible factor 1, alpha (Hif1-α).^23,25^ Furthermore, WAT health is also linked to thermogenesis,^26^ a process driven by brown adipose tissue (BAT) and the emergence of beige adipocytes within WAT, known as “beiging”. Increasing the prevalence of thermogenic adipocytes leads to higher energy expenditure and a lean, metabolically healthy phenotype.^22,27^

Procollagen endopeptidase enhancer protein 2 (Pcpe2) is found in the extracellular matrix (ECM) and is a paralog of Pcpe1, both of which are highly expressed in adipose tissue PCs.^17,28,29^ Although these glycoproteins are better known for their participation in the maturation of fibrillar collagens,^30,31^ they also appear to function in other capacities.^32^ Previous studies have established that the enhancing activity of Pcpe1 is regulated by domains called **C**omplement C1r/C1s, **U**EGF, **B**MP-1, or **CUB**s, which facilitate optimal substrate presentation to bone morphogenetic protein 1 (BMP1), an enzyme that aids in collagen maturation.^33^ Pcpe1 binds specifically to the C-propeptide region of procollagen, while it has been reported that Pcpe2 interacts with both BMP1 and the C-terminal propeptide of procollagens.^31,34^ Owing to their substantial structural homology,^32,35^ Pcpe1 and 2 were initially thought to perform similar biological roles.^35^ Initially, structural features based on their glycosylation were noted￼ ;^35^however, it was never fully investigated, and now accumulating evidence indicates that, despite their amino acid sequence similarity, Pcpe1 and 2 do exert distinct, and in some cases opposing, effects on procollagen processing and tissue remodeling.^31^ Pcpe2, in contrast to Pcpe1, is less well studied and was identified as a collagen-binding protein mapped to chromosome 3 in humans and to chromosome 9 in mice, with approximately 45.6% structural similarity to Pcpe1.^32^ Interestingly, the amino acid sequence of Pcpe2 in humans is quite similar to that in mice, with ∼89.1% amino acid homology, and remains highly conserved down the phylogenetic tree, with 81.5% amino acid homology between humans and coelacanth.^32^ On the other hand, the amino acid divergence of Pcpe1 between humans and coelacanths is ∼51.1%.^32^ Overall, these two ECM glycoproteins have been relatively understudied, especially with regard to their physiological roles in cardiometabolic disease progression.

Looking beyond Pcpe1 and 2’s structural similarities, several recent functional studies using genetically engineered mouse models have been reported. For example, in global *Pcolce*^KO^ mice, abnormal collagen fibrils were observed in both mineralized and nonmineralized tissues, with 100% of tendon collagen fibrils exhibiting abnormal morphologies.^36^ Another group studied global *Pcolce1*^KO^ mice and reported significantly reduced liver fibrosis, body weight, and collagen I levels, but no reduction in the progression of nonalcoholic steatohepatitis.^37^ This same group also showed that global *Pcolce1*^KO^ (gene name for Pcpe1) mice had significantly lower cardiac ejection fraction, with no changes in cardiac collagen content or wound healing.^38^ These authors concluded that Pcpe1 deficiency in mice did not affect cardiac collagen content or cardiac function under non-stressed conditions.^37^ However, in other reports investigating both global and tissue-specific Pcpe1 knockout mice,^39,40^ research showed that Pcpe1 acts as a BAT-derived cytokine that promotes liver fibrosis and induces nonalcoholic steatohepatitis in mice.^39,40^ The authors concluded that mice lacking Pcpe1 showed markedly ameliorated liver fibrosis, suggesting that Pcpe1 functions beyond its established role in BMP1 enhancement of collagen processing, and more like a secreted signaling molecule with cytokine-like properties that facilitate inter-organ communication.

Investigative studies involving Pcpe2 have been far fewer. The first *Pcolce2* global knockout mouse study showed normal thymic epithelial cell differentiation,^41^ while in another report, *Pcolce2^KO^* (gene name for Pcpe 2) mice exhibited myocardial stiffness following transverse aortic constriction, suggesting that Pcpe2 contributed to both myocardial compliance and collagen deposition in the mouse heart.^42^ In lipid-related studies, Pcpe2 was reported to enhance BMP1-mediated cleavage of pro-apoA-I, a crucial step in the production of mature plasma apoA-I, the main protein constituent of high-density lipoprotein (HDL).^43,44^ Consistent with this idea, this same group showed an approximately twofold higher plasma concentration of enlarged HDL particles in *Pcolce2^KO^* mice, suggesting that the lack of Pcpe2’s enhancement of BMP1-mediated pro-apoA-I processing in mice reduced cholesterol efflux capacity.^45,46^ Based on the results of these studies, our lab further investigated Pcpe2’s role in lipid metabolism. Pollard et al^47^, crossed *Pcolce2^KO^* mice with low-density lipoprotein receptor (LDLR) knockout mice and observed enhanced progression of diet-induced atherosclerosis in double knockout mice compared to controls. These data suggested that loss of Pcpe2 led to dysfunctional removal of plasma HDL via the scavenger receptor class B type I (SR-BI) in the liver.^47,48^ In other studies involving Pcpe2 and lipid metabolism, a study in type 2 diabetic (T2D) patients identified the *PCOLCE2* gene among the top differentially expressed genes (DEGs) in sorted β-cells.^49^ In a functional follow-up, results indicated that the reduction in Pcpe2 expression was associated with reduced mitochondrial activity and contributed to impaired insulin secretion.^49^ In another study, using differentiated adipocytes from mice, Pcpe2 expression was found to alter lipid storage.^50^ Using global *Pcolce2^KO^*mice, the data showed reduced SR-BI-mediated HDL uptake in mature adipocytes, thereby compromising adipocyte lipid storage. Taken together, these findings illustrate that Pcpe2 is a multifunctional protein with roles extending beyond ECM regulation to include lipid metabolism and that it appears to impact cellular plasticity, tissue regeneration, and adipose tissue homeostasis.

In the current study, we further elucidate the cell-autonomous role of Pcpe2 in adipose tissue remodeling using mice with a specific adipocyte-specific deletion of Pcpe2 (TgAdip^+^Pcpe2^KO^). These studies demonstrate that the loss of Pcpe2 specifically in adipose tissue profoundly improves systemic metabolic health and fat storage, with WD-fed TgAdip^+^Pcpe2^KO^ mice exhibiting significantly lower body weight and adipose tissue mass, coupled with improved glucose homeostasis and increased energy expenditure. These metabolic improvements are associated with a dramatic suppression of the pathological hallmarks of obesity, including chronic inflammation, macrophage infiltration, and a shift in PC populations toward adipogenic precursors, with a significant increase in thermogenic capacity. Our studies also point to a functional role for a mucin-like linker domain, which also distinguishes Pcpe2 from Pcpe1. Collectively, these findings establish Pcpe2 as a critical, novel regulator of adipose tissue health and energy expenditure, offering a promising molecular target for the treatment of obesity and related cardiometabolic disorders.

## Materials and Methods

### Animals and Diet

All animals were reviewed and approved by the Institutional Animal Care and Use Committee at the Medical College of Wisconsin. Conditional Pcpe2 knockout and over-expressor mice were generated in collaboration with Cyagen Biosciences. For Tissue-specific Pcpe2 knockout, using C57BL/6NTac background, we generated loxP targeted Pcpe2 mice, where exon 3 was flanked by loxP sites. This was done with the help of the Cyagen Biosciences, Santa Clara, USA. A codon-optimized construct was designed to flank exon 3 of the *Pcolce2* gene with loxP sites and added to AAV vector that expresses Cas9 and gRNAs to target double-strand breaks around exon 3 of the *Pcolce2* locus. AAV particles were then incubated with zygotes to induce breaks and homology-directed repair. Positive mice were obtained, and the resulting *Pcolce2-loxP* mice were bred with Adiponectin-Cre mice (strain 028020) or Pdgfra-Cre mice (strain 013148), acquired from Jackson Laboratory, to allow adipose-specific and precursor cell-specific deletion of Pcolce2, respectively.

Pcpe2 overexpressing (Pcpe2^Ox^) mice were generated using a conditional ROSA26 knock-in approach, in collaboration with Cyagen Biosciences. The targeting construct was designed so that the mouse Pcpe2 coding sequence followed by GGGGS (linker) and 5-hemagglutinin (HA) tag was inserted into the ubiquitously active ROSA26 safe-harbor locus under the control of a strong CAG promoter. Transcriptional STOP cassette flanked by loxP sites was placed upstream of the Pcpe2 sequence. In the absence of Cre, the STOP cassette prevented expression, keeping Pcpe2 silent in the knock-in line. When crossed with Cre-expressing driver lines (Adiponetin-Cre and/or Pdgfra-Cre), Cre recombinase excised the loxP-flanked STOP cassette, thereby bringing the Pcpe2 coding sequence under the direct control of the CAG promoter. Mouse genotypes from tail biopsies were determined with specific probes designed for each gene (Transnetyx, Cordova, TN). Mice were fed a standard (chow) (SD) diet and a Western diet (WD) (Teklad^TM^ TD.88137) for 20-30 weeks. The Western diet contained 0.2% cholesterol and 21% total fat, contributing to 42% kcal from fat.

Adiponectin^Cre/+^ RosaDTA^fl/+^mice were generated by crossing Adiponectin^Cre^ mice (The Jackson Laboratory, Bar Harbor, ME, USA: 028020) with lox-stop-lox-Rosa26 diphtheria toxin (DTA) mice (The Jackson Laboratory: 010527: RRID: IMSR_JAX:010527), resulting in the ablation of mature adipocytes and consequent lipoatrophy in Adiponectin^Cre/+^ RosaDTA^f/+^ offspring. These lipoatrophic mice, which constitutively lack brown and white adipose tissue, and their littermate controls were maintained at thermoneutrality (30°C) from birth on a 12-h light/dark cycle.

Throughout the terms VAT, SAT, and BAT will be used exclusively to indicate the following mouse anatomical depots: VAT refers to peri-gonadal fat; SAT refers to inguinal fat; and BAT refers to interscapular fat.

### Single-Cell RNA Sequencing Information from Publicly Available Portals

Tissue distribution and cell type-specific expression of PCPE2 in human and mouse adipose tissues were analyzed using publicly available-cell RNA sequencing (scRNA-seq) datasets. Mouse and human adipose scRNA-seq data were obtained from the Single Cell Portal at the Broad Institute of MIT and Harvard.^51^ The data are derived from the comprehensive WAT atlas, which encompasses multiple adipose depots and cell populations, including adipocytes, adipose stem and progenitor cells (ASPCs), endothelial cells, and immune cells.^17^ UMAP visualizations of human scRNA-seq data were obtained from the Adipose Tissue Knowledge Portal,^28^ which compiles and harmonizes datasets as described in three publications that collectively provide a cross-species view of PCPE2/Pcpe2 expression across major adipose tissue cell populations and metabolic states.^29,52,53^

### Glucose Tolerance Test

For the glucose tolerance test (GTT), mice were fasted for 3-4 hours before receiving an intraperitoneal injection of 2.5 g/kg body weight of glucose. Blood samples were collected from the tail vein at 0, 20, 40, 60, 80, 100, and 120 minutes post-injection, and glucose concentrations were determined using a Contour Next EZ glucometer. Plasma insulin concentrations were quantified using a Rat/Mouse Insulin ELISA Kit (Millipore).

### Plasma Cholesterol and Triglyceride

Cholesterol levels were measured using the Amplex® Red Cholesterol Assay Kit (Thermo Fisher Scientific) following the manufacturer’s protocol. Briefly, samples or cholesterol standards were incubated with Amplex® Red reagent, horseradish peroxidase, cholesterol oxidase, and cholesterol esterase. The enzymatic reaction generates resorufin, which was quantified fluorometrically (Ex 560 nm/Em 590 nm) using a microplate reader. Plasma triglyceride levels were quantified using the Triglyceride Assay Kit (FUJIFILM/Cayman) according to the manufacturer’s instructions. Briefly, plasma samples were diluted in assay buffer and incubated with the enzymatic reagent containing lipoprotein lipase, glycerol kinase, glycerol phosphate oxidase, and peroxidase. The enzymatic reaction produces a chromogenic compound proportional to triglyceride concentration, which was measured spectrophotometrically at 600 nm using a microplate reader.

### Fast Protein Liquid Chromatography

FPLC was conducted at 4°C using AKTA purifier 10 with two columns of Superose 6 increase 10/300 GL (Cytiva, Cat# 29091596) in tandem. 300 μl (1.5 times of the volume of the sample loop) of pooled plasma was injected. Tris-buffered saline (25 mM Tris, 150 mM NaCl, 2 mM EDTA, pH = 7.4) was used as the running buffer. The eluant was collected in 1 mL fractions from fractions 12 ml to 47 ml.

### Isolation of Mature Adipocytes and Stromal Vascular Fraction from Adipose Tissue

SVF was prepared from freshly isolated mouse adipose tissue as described previously.^54,55^ Adipose depots were dissected under sterile conditions and placed into a beaker of sterile PBS. Filling a 1.5 ml tube with adipose tissue, the tissue was then minced using spring scissors until no visible tissue fragments remained. The minced material was transferred into a 50 ml Falcon tube containing 10 ml of freshly made digestion buffer consisting of 1 mg/ml of collagenase D (Roche #64311324) in Hank’s Balanced Salt Solution with Ca^2+^ and Mg^2+^ (Gibco#14025-092), and 1.5% bovine serum albumin (Sigma Aldrich #A9647). The suspension was briefly vortexed to disperse tissue fragments and incubated at 37°C in a shaking incubator for 60 minutes (VAT) and 75 minutes (SAT and BAT) to ensure complete enzymatic digestion. After digestion, the cell suspension was filtered through a 100 µm cell strainer to remove residual undigested material. The filtrate was centrifuged at 600g for 5 minutes at room temperature. The floating mature adipocyte layer was carefully removed and used for downstream applications.

### Differentiation of Stromal Vascular Fraction Cells

The freshly isolated SVF pellet was resuspended in red blood cell lysing buffer (Sigma #64311324) and incubated for 1 minute at room temperature with gentle mixing to eliminate erythrocyte contamination. Lysis was quenched by adding Growth Medium composed of DMEM/F12 supplemented with GlutaMAX (Gibco #10565018), 10% fetal bovine serum (FBS) (Sigma F0926), and 1% penicillin-streptomycin (Gibco 15140122). The suspension was passed through a 40 µm cell strainer and centrifuged again at 600g for 5 minutes at 4°C. The resulting SVF pellet was either used in other downstream applications or resuspended in pre-warmed Growth Medium and plated onto a 60 mm collagen-coated culture dish (Corning #354401). Cells were maintained at 37°C in a humidified atmosphere containing 5% CO₂. Growth Medium was replaced the following day to remove non-adherent cells and debris. After a total of 2 days in culture, the confluent cells were treated with TrypLE Express (Gibco #12605), counted, then plated at 20,000 cells per 96-well plate or 100,000 cells per 24-well plate. In our studies, one 60 mm dish referenced above is derived from the equivalent of 4 SAT pads from 2 male mice fed a standard diet and between the ages of 6-12 weeks. Two days following replating, the cells were treated with Induction Medium, consisting of Growth Medium with 5 ug/ul Insulin (Sigma #19278), 1 uM Dexamethasone (Sigma #D442), 500 uM IBMX (Sigma #15879), and 1 uM Rosiglitazone (Sigma #R2408) for 2 days. After induction of differentiation, the cells are maintained in Maintenance Medium, consisting of Growth Medium supplemented with 5 ug/ml insulin for up to 4 days.

### FACS Analysis and FAP and APC Sorting

FACS analysis was performed on freshly isolated SVF cells after incubating with fluorophore-conjugated antibodies against CD45 (Biolegend, 103132), CD31 (Biolegend, 1024220), CD140b (Pdgfrb) (Biolegend, 136006), Ly6C (Biolegend, 128016), and CD9 (Biolegend, 124808). The gating strategy for distinguishing FAPs and APCs was selected as previously describe^18^. Dead cells and debris were excluded based on forward- and side-scatter properties. To eliminate immune and endothelial cell contaminants, cells positive for CD45 and CD31 were first gated. The CD45⁻CD31⁻ population was then analyzed for CD140b^+^ (Pdgfrb) expression to identify precursor cells. Within this CD140b⁺ (Pdgfrb⁺) population, further discrimination based on Ly6C and CD9 expression defined two distinct subsets: fibroadipogenic precursor cells (FAPs; Pdgfrb⁺Ly6C⁺CD9⁺) and adipogenic precursor cells (APCs; Pdgfrb⁺Ly6C⁻CD9⁻). FAPs and APCs were sorted using a BD-FACSAria II cell sorter (BD Biosciences). Sorted populations were collected in growth medium containing 10% FBS and maintained on ice until used for cell culture or their RNA extracted using RNAqueous-Micro Kit (Invitrogen #AM1931).

### Adipogenic Index

Freshly isolated SVF cells were seeded into 96-well black plates (Greiner Bio-One #655090) and maintained at 37°C in a humidified CO₂ incubator. Light microscopy images were taken daily to assess lipid droplet formation. To calculate the adipogenic index,^15^ cells were incubated with Bodipy 493/503 (Invitrogen #191867) for 30 minutes and counterstained with Hoechst 33342 nuclear stain (Thermo Fisher Scientific #62249) for 30 minutes at 37°C. After washing each well three times with 1× PBS, fluorescence images were acquired. Images and quantitative determination of total Bodipy lipid staining area per cell were performed using the Keyence BZ-X800 Microscope.

### Seahorse Analysis

SVF cells from the SAT of standard diet-fed male mice were obtained as described and plated at 20,000 cells per 96-well onto collagen-coated Seahorse XFe96 cell culture microplates (Agilent Technologies #103794). Cells were differentiated as described above. The mitochondrial respiration and glycolytic function were assessed by measuring oxygen consumption rate (OCR) and extracellular acidification rate (ECAR), respectively using the Seahorse XFe96 Extracellular Flux Analyzer in the MCW Redox and Bioenergetics Core (RRID:SCR_027984) within the Wisconsin Cancer Center Translational Metabolomics Shared Resource, following the manufacturer’s recommended protocols. Data were normalized by cell number per well using Hoechst nuclear staining and imaged using a Keyence BZ-X800 Microscope. Data were analyzed using Seahorse Wave software.

### Lentiviral Transduction

Wild-type human and mouse PCPE2/Pcpe2 plasmids were obtained from OriGene (Rockville, MD) and modified for His-tag protein expression (TOP Gene Technologies Inc), as previously described^50^. The wild-type human PCPE2 plasmid was further modified to replace each linker region’s ninth threonine with alanine to prevent potential O-linked glycosylation,^32^ creating a non-O-glycosylated (NG-PCPE2) construct.^50^ Lentiviral particles were produced at the Versiti Blood Research Institute Core Facility (RRID:SCR_025503). These particles were added at an MOI of 10 SVF cells, prepared as described above, which were originally seeded into 96-well plates at a density of 20,000 cells per 96-well (Greiner Bio-One #655090). During plating/transduction, cells were treated with polybrene at a final concentration of 0.5 µg/µl to enhance infection efficiency. The cells were maintained at 37°C in a humidified incubator with 5% CO₂ and then differentiated into adipocytes as described above.

### DiI-VLDL Uptake Assay

Differentiated SVF cells were prepared as described above. Following differentiation, the maintenance medium was removed, and the cells were washed three times with 1× PBS. Cells were then incubated in serum-free and phenol red-free DMEM (Gibco #31053-028), and baseline fluorescence was measured using a CLARIOstar plate reader (excitation 541–15 nm; emission 590–20 nm) to obtain background plate readings. Cells were subsequently incubated with DiI-labeled VLDL (Kalen Biomedical #770130-9) at a final concentration of 10 µg/ml in serum-free and phenol red-free DMEM for 90 minutes at 37°C in a humidified incubator with 5% CO₂. Following DiI-VLDL uptake, the media was removed from each well and cells were washed three times with 1× PBS and replenished with 100 µl phenol red–free DMEM per well. Fluorescence was measured again using the CLARIOstar settings (excitation 541-15 nm; emission 590-20 nm). Background fluorescence values were subtracted from the DiI-VLDL signal. To normalize for cell number, each well was treated with 1× CyQUANT reagent (Thermo Fisher Scientific, C35011) for 1 hour at 37°C in 5% CO₂ following the manufacturer’s recommendations. Fluorescence was measured (excitation 483–14 nm; emission 530-30 nm), and the DiI-VLDL fluorescence uptake per well was normalized to cell number.

### RNA Isolation and Quantitative RT-PCR

Total RNA was isolated from cells using the RNeasy Mini Kit (Qiagen #74104) for SVF and RNeasy Lipid Tissue Mini Kit (Qiagen #74804) for adipocytes, following the manufacturer’s instructions. Concentrations of isolated RNA were determined using a NanoDrop spectrophotometer. cDNA was synthesized using the iScript cDNA synthesis kit (Bio-Rad #1708891). Quantitative real-time PCR was performed with iTaq Universal SYBR Green Supermix (Bio-Rad#1725121) on a Bio-Rad CFX384/C1000 detection system, using 18S rRNA as an internal control for normalization. All primers were obtained from Integrated DNA Technologies (Coralville, IA). Relative gene expression was analyzed using the ΔΔCt method.

### Tissue Harvest, Fixation, and Histological Staining

At the time of necropsy, liver, VAT. SAT and BAT depots were removed and weighed to assess tissue mass. Tissues were immersed in 4% paraformaldehyde-PBS fixative for 48 hours (ChemCruz #sc-281692), then stored in 70% ethanol before being processed and embedded in paraffin. Tissue sections of 7 µm were stained with hematoxylin and eosin (H&E) or Masson’s trichrome. For immunofluorescent staining, sections were incubated with antibodies to CD68 (Cell Signaling #97778), Perilipin (Invitrogen #MA5-27861), and UCP1 (Abcam #10983) by the Pathology Histology Core Facility at MCW, (RRID:SCR_028288). High-resolution images were acquired using a Keyence BZ-X800 Microscope.

### Mouse Metabolic Phenotyping

Comprehensive metabolic phenotyping was carried out using a 16-chamber Promethion metabolic monitoring system (Sable Systems International) in the MCW Comprehensive Rodent Metabolic Phenotyping Core (RRID:SCR 028283).^56,57^ Body composition, including fat, lean and fat-free mass, was determined using time-domain nuclear magnetic resonance (TD-NMR; Bruker LF110). Data were analyzed using CalR software.^58^

### Fat Transplantation

VAT depots collected from donor TgAd^+^Pcpe2^ox^ mice or WT littermates, aged 10-14 weeks, were transplanted into the scruff of 7-10 week old lipoatrophic Adiponectin^Cre/+^ RosaDTA^fl/+^recipient male mice to generate recipients whose adipose tissue was the sole carrier of Pcpe2^ox^ genotype (or control) in the body. The surgeries were conducted as previously described^59,60^ with a few modifications. After inducing anesthesia by continuous inhalation of 1-2% isoflurane, with mice placed on a heating pad in the prone position, a 1-2 mm incision was made in the dorsal scruff. Three small pockets were made bilaterally, and one graft (150-300 mg) was implanted into each pocket, for a total of six fat pads per mouse, all confined to the subcutaneous dorsal space. Incisions were closed with 5-0 nylon sutures. The mice were given a subcutaneous injection of extended-release buprenorphine (1 mg/kg) and placed overnight in an infant incubator warmed to 27°C. Mice were housed for 10 weeks, with weekly weight measurements and occasional submandibular bleeds to assess circulating triglycerides, cholesterol, and glucose.

### Mass Spectrometry Sample Preparation

Recombinant wild-type human PCPE2 and NG-PCPE2 were reconstituted in 50 mM ammonium bicarbonate. PCPE2 was first digested with SmE, which cleaves immediately N-terminal to Ser or Thr residues bearing mucin-type O-glycans, for 3 h at an enzyme-to-substrate ratio of 1:10. SmE was expressed in-house as previously described.^61^ Both human PCPE2 and NG-PCPE2 samples were reduced with 1 mM dithiothreitol (Sigma, D0632) at 65 °C for 20 min and alkylated with iodoacetamide (Sigma, I1149) at a final concentration of approximately 1.6 mM for 15 min at room temperature in the dark. Reduced and alkylated samples were subsequently digested with sequencing-grade modified trypsin (Promega, V511A) at a 1:50 enzyme-to-substrate ratio overnight at 37°C, generating glycopeptides spanning the mucin-like linker domain with both O-glycoprotease- and trypsin-derived termini. Reactions were quenched by adding 1 µL of formic acid (Thermo Scientific, 85178) and diluted to a volume of 200 µL prior to desalting. Desalting was performed using 10 mg Strata-X 33 µm polymeric reversed phase SPE columns (Phenomenex, 8B-S100-AAK). Each column was activated using 500 µL acetonitrile (ACN) (Honeywell, LC015) followed by 500 µL 0.1% formic acid, 500 µL 0.1% formic acid in 40% ACN, and equilibration with two additions of 500 µL 0.1% formic acid. After equilibration, the samples were added to the column and rinsed twice with 200 µL 0.1% formic acid. The columns were transferred to a 1.5 mL tube for elution by two additions of 150 µL 0.1% formic acid in 40% ACN. The eluent was then dried using a vacuum concentrator (LabConco) prior to reconstitution in 10 µL of 0.1% formic acid.

### Mass Spectrometry Data Acquisition

Samples were analyzed by online nanoflow liquid chromatography-tandem mass spectrometry using an Orbitrap Eclipse Tribrid mass spectrometer (Thermo Fisher Scientific) coupled to a Dionex UltiMate 3000 HPLC (Thermo Fisher Scientific). For each analysis, 4 ul was injected onto an Acclaim PepMap 100 column packed with 2 cm of 5 µm C18 material (Thermo Fisher, 164564) using 0.1% formic acid in water (solvent A). Peptides were then separated on a 15 cm PepMap RSLC EASYSpray C18 column packed with 2 µm C18 material (Thermo Fisher, ES904) using a gradient from 035% solvent B (0.1% formic acid with 80% acetonitrile) in 60 min. Full scan MS1 spectra were collected at a resolution of 60,000, an automatic gain control target of 3e5, and a mass range from m/z 300 to 1500. Dynamic exclusion was enabled with a repeat count of 2, repeat duration of 7 s, and exclusion duration of 7 s. Only charge states 2 to 6 were selected for fragmentation. MS2s were generated at top speed for 3 s. Higher-energy collisional dissociation (HCD) was performed on all selected precursor masses with the following parameters: isolation window of 2 m/z, 29% normalized collision energy, orbitrap detection (resolution of 7,500), maximum inject time of 50 ms, and a standard automatic gain control target. An additional electron transfer dissociation (ETD) fragmentation of the same precursor was triggered if 1) the precursor mass was between m/z 300 to 1500 and 2) 3 of 8 HexNAc or NeuAc fingerprint ions (126.055, 138.055, 144.07, 168.065, 186.076, 204.086, 274.092, and 292.103) were present at m/z ± 0.1 and greater than 5% relative intensity. Two files were collected for each sample: the first collected an ETD scan with supplemental energy (EThcD) while the second method collected a scan without supplemental energy. Both used charge calibrated ETD reaction times, 100 ms maximum injection time, and standard injection targets. EThcD parameters were as follows: Orbitrap detection (resolution 7,500), calibrated charge-dependent ETD times, 15% nCE for HCD, maximum inject time of 150 ms, and a standard precursor injection target. For the second file, dependent scans were only triggered for precursors below m/z 1000, and data were collected in the ion trap using a normal scan rate.

### Mass Spectrometry Data Analysis

Raw files were searched using Byonic (version 4.5.2, Protein Metrics, Inc.) against a curated database including the human PCPE2 sequence and a mutated human PCPE2 sequence changing each of the Tyr residues in the mucin-like linker domain to Ala (NG-PCPE2). For all samples, we used the default O-glycan database containing 9 common structures. Files were searched with semi-specific cleavage N-terminal to Ser and Thr and six allowed missed cleavages. Samples treated with trypsin were searched with the same parameters but also allowed cleavage C-terminal to Arg or Lys. Mass tolerance was set to 10 ppm for MS1’s and 20 ppm for MS2’s. Met oxidation was set as a variable modification and carbamidomethyl Cys was set as a fixed modification. From the Byonic search results, glycopeptides were filtered to a score of >200 and a logprob of >2. From the remaining list of glycopeptides, the extracted ion chromatograms, full mass spectra (MS1s), and fragmentation spectra (MS2s) were investigated in XCalibur QualBrowser (Thermo) to generate a list of true-positive glycopeptides, as reported in **Supplemental Table 1**. Each reported glycopeptide listed in **Supplemental Table 1** was manually validated from the filtered list of Byonic’s reported peptides (score>200 and logprob >2) according to the following steps: The MS1 was first used to confirm the precursor mass and chosen isotope was correct. This also allowed us to identify any co-isolated species that could interfere with the MS2s and/or explain unassigned peaks. The HCD and EThcD fragmentation spectra were then investigated to identify sufficient coverage to make a sequence assignment. When possible, multiple MS2 scans were averaged to obtain a stronger spectrum. For HCD, an initial glycopeptide identification was confirmed if the presence of the precursor mass without a glycan present (i.e., Y0), along with coverage of b and y ions without glycosylation. For longer peptides, we required the presence of Y0 and fragments that were expected to be abundant (e.g., N-terminally to Pro, C-terminally to Asp). When the peptide contained a Pro at the C-terminus, the bn-1 was considered sufficient. We then used electron-based fragmentation MS2 spectra for localization. Here, all plausible localizations were considered, regardless of search result output. We confirmed the presence of fragment ions in ETD or EThcD that were between potential glycosylation sites, if sufficient c/z ions were present then a glycan mass was considered localized. For glycopeptide manual validation, extracted ion chromatograms are evaluated at the MS1 level to determine the charge and m/z of the highest abundance precursor species. To determine individual glycopeptide abundances, we first generated extracted ion chromatograms of monoisotopic masses and calculated areas under the curve (AUCs). To avoid bias toward smaller glycan structures while minimizing the inconvenience of typing multiple isotope masses into XCalibur QualBrowser (Thermo) for each peptide, abundances were calculated using a variable number of isotopes. Only the 12C peak was used for anything under 1600 Da, the 13C was also included up to 2400 Da, and three isotopes were collected over 2400 Da. These values were chosen based on the predicted isotopic distributions for given intact peptide values to allow the majority of the signal to be incorporated. All charge states were included when determining the abundance of a given glycoform. Mass spectrometry data files and raw search output can be found on PRIDE with identifier PXD064788.

### Data Availability Statement

All mass spectrometry data and search results acquired for this manuscript have been deposited on the PRIDE repository. Reviewers can access raw data with the following login information: Project accession: PXD071066, Token: iD7BJE9KjF3W. Alternatively, the dataset can be accessed by logging in to the PRIDE website using the following account details: Username: reviewer Password: f6AiaQz4kfK3

### Statistics

Data are expressed as mean ± standard deviation. Statistical analyses were performed using GraphPad Prism version 11 (GraphPad Software, San Diego, CA). For comparisons between two groups, Student’s t-test was applied to data that met normality assumptions using the Shapiro-Wilk test, whereas the Mann–Whitney U test was used for non-normally distributed data. For experiments involving more than two groups, statistical significance was determined by one-way or two-way ANOVA, as appropriate, followed by Tukey’s multiple comparisons test for post-hoc analysis.

### Human Study Design and Methods

The adipose tissue samples analyzed in Studies 1 and 2 were obtained from a subset of participants who completed studies evaluating cardiometabolic characteristics of people with metabolically healthy and unhealthy obesity^62^ and the effects of weight loss on metabolic function.^63^ Both studies were approved by the Human Research Protection Office at Washington University School of Medicine in St. Louis, MO and registered in ClinicalTrials.gov (NCT02706262 and NCT02207777, respectively). Written informed consent was obtained from all participants before they started the studies, which were conducted in the Clinical Translational Research Unit (CTRU) at Washington University School of Medicine in St. Louis, MO.

Study 1 involved 50 men and women who were metabolically healthy lean (n=14; 7 men and 7 women), metabolically healthy obese (n=18; 1 man and 17 women) or metabolically unhealthy obese (n=18; 2 men and 16 women). The following inclusion criteria were required for each cohort: i) metabolically healthy lean (MHL) defined as BMI 18.5-24.9 kg/m^2^, fasting plasma glucose concentration <100 mg/dL, 2-hr oral glucose tolerance test (OGTT) plasma glucose concentration ≤140 mg/dL, HbA1c ≤5.6% and normal whole-body insulin sensitivity, defined as the glucose infusion rate (GIR) per kg fat-free mass (FFM) divided by the plasma insulin concentration (GIR/Insulin) during the final 20 minutes of the hyperinsulinemic-euglycemic clamp procedure >40 (µg/kg FFM/min)/(µU/mL); ii) metabolically healthy obese (MHO) had BMI 30-49.9 kg/m^2^, fasting plasma glucose concentration <100 mg/dL, 2-hr OGTT plasma glucose concentration ≤140 mg/dL, HbA1c ≤5.6% and normal whole-body insulin sensitivity; and iii) Metabolically unhealthy obese (MUO) had BMI 30-49.9 kg/m^2^, HbA1c 5.7%-6.4% or fasting plasma glucose concentration 100-125 mg/dL or 2-hr OGTT plasma glucose concentration 140-199 mg/dL and impaired whole-body insulin sensitivity, defined as a GIR/Insulin ≤40 (µg/kg FFM/min)/(µU/mL). Potential participants who had a history of diabetes or liver disease other than metabolic dysfunction-associated steatotic liver disease (MASLD), were taking medications that can affect metabolism, had metal implants that precluded magnetic resonance imaging or consumed excessive amounts of alcohol (>21 units of alcohol per week for men and >14 units of alcohol per week for women) were excluded. Study 2 involved 15 people with obesity and type 2 diabetes (T2D) (age 53±3 years; 4 men and 11 women). Participants were included if they achieved marked (>15%) weight loss, induced by either Roux-en-Y gastric bypass surgery (n=6) or low-calorie diet therapy (n=9). This weight loss target was chosen because it is associated with a high rate of remission of T2D.^64^ Potential participants who had evidence of significant organ system dysfunction or disease other than MASLD and T2D, previous intestinal resection, or consumed excessive amounts of alcohol (>21 oz of alcohol per week for men and >14 oz of alcohol per week for women) were excluded.

### Human Adipose Tissue Biopsies

Subcutaneous abdominal adipose tissue (SAAT) was obtained from the periumbilical after an overnight fast in both Studies 1 and 2. After anesthetizing the skin by percutaneous injection of 1% lidocaine, a small skin incision (∼0.5 cm) was made and ∼0.5 grams of adipose tissue was aspirated through a 4-mm liposuction cannula (Tulip Medical Products, San Diego, CA) connected to a 60 cc syringe. These samples were immediately rinsed in ice-cold saline and frozen in liquid nitrogen before being stored at -80°C until processed for RNA sequencing. The RNA-sequencing data have been deposited in the NCBI Gene Expression Omnibus database (GSE244121 for Study 1 and GSE156906 for Study 2).

## Results

### Pcpe2 Expression Correlates with Adipose Tissue Mass

We next explored the tissue expression profile of Pcpe2, a homolog of Pcpe1.^32,65^ In human adipose tissue, *PCOLCE2* (human gene name) mRNA and *Pcolce2* in the mouse were compared by searching multiple online databases. We found that both are highly expressed in human and mouse VAT, SAT, and BAT.^66^ At the cellular level, human and mouse *Pcolce2* mRNA are both highly expressed in mature adipocytes and PCs, specifically, adipocyte stem precursor cells (ASPCs) and/or fibroadipogenic precursors (FAPs) **(Figures 1A and G**). These data indicate that in humans, Pcpe2 expression is somewhat higher in mature adipocytes than PCs, whereas in mice, expression is higher in PCs than in mature adipocytes. Comparatively, *Pcolce1* mRNA expression appeared predominantly in PCs over mature adipocytes **(Supplemental Figure 1A and F).** Both glycoproteins showed minimal expression in other cells found in adipose tissue, including immune and endothelial populations **(Figure 1B and Supplemental Figure 1B)**. Interestingly, time-course analyses showed that Pcpe2 mRNA expression (and Pcpe2 proteomic expression) was elevated in SVF PCs, which steadily increased during differentiation into mature adipocytes **(Figure 1C).** Conversely, Pcpe1 mRNA expression remained relatively constant over time and decreased slightly during differentiation and maturation **(Supplemental Figure 1C).**

**Figure 1.**
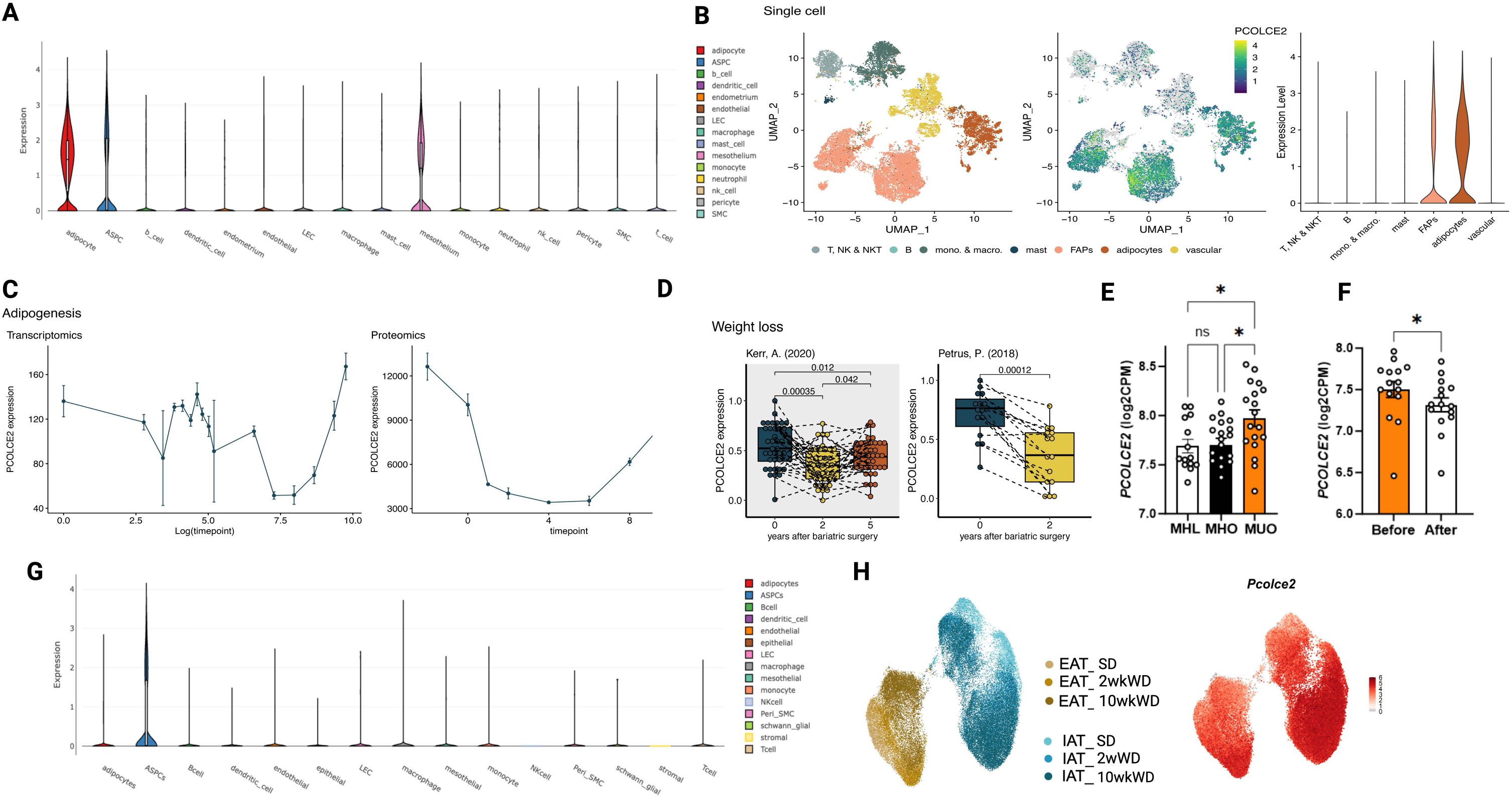
Expression of *PCOLCE2 I Pcolce2* and Adipose Tissue Mass in Humans and Mice. **(A)** *PCOLCE2* relative expression among various cell types as determined by single-cell RNA sequencing (scRNA-seq) of human white adipose tissue (WAT) **(B)** UMAP of human WAT *PCOLCE2* expression in fibroinflammatory adipocyte precursors (FAPs) and mature adipocytes relative to various immune cells **(C) (Left panel)** *PCOLCE2* mRNA expression and **(Right panel)** PCPE2 protein expression expressed during adipocyte differentiation **(D)** Human *PCOLCE2* mRNA expression from subcutaneous adipose tissue (SAT) collected before and after weight loss. Studies involved obese individuals who underwent bariatric surgery, with follow-up assessments conducted **(Left panel)** two and five years post-surgery as reported by Kerr et al., 2020 or **(Right panel)** two years post-surgery in the study reported by Petrus et al., 2018 **(E)** *PCOLCE2* mRNA levels from human patients designated metabolically healthy lean (MHL; n=19), metabolically healthy obese (MHO; n=18) and metabolically unhealthy obese (MUO; n=14). Data are presented as mean ± SEM. Statistical significance was determined by one-way ANOVA with *p < 0.05 **(F)** *PCOLCE2 mRNA* expression before and after 18% weight loss (n = 15). The weight loss group represents 9 people who lost weight by consuming a low-calorie diet and 6 that had Roux-en-Y gastric bypass surgery **(G)** scRNA-seq of mouse WAT showing *Pcolce2* expression among various cell types **(H) (Left Panel)** UMAP from public scRNA-seq dataset (Nahmgoong, Hahn, et al., 2022) showing mouse adipose stem cells from epididymal adipose tissue (EAT) and inguinal adipose tissue (IAT) from three dietary conditions: normal standard diet (SD), 2-week Western diet (2wWD), and 10-week Western diet (10wWD). Colors indicate tissue origin and dietary condition. **(Right panel)** UMAP of *Pcolce2* gene expression across dietary conditions. **Note:** *PCOLCE2/Pcolce2* scRNA-seq data of human **(A)** and mouse **(G)** were obtained from the single cell portal (Broad Institute of MIT and Harvard; Tarhak et al., 2025), based on datasets from Emont et al., 2022. Adipogenesis data and UMAP of human scRNA-seq **(B and C)** were acquired from from the adipose tissue Knowledge portal (Zhong et al., 2025), which includes data from Massier et al., 2023; Hinte et al., 2024; and Reinisch et al., 2024.

Portal-derived data also showed significant associations between SAT Pcpe2 mRNA abundance and body mass, but not between Pcpe1 mRNA abundance and body mass. Two independent studies involving obese patients **(Figure 1D and Supplemental Figure 1D)** suggest that Pcpe2 mRNA levels are significantly reduced with fat loss 2 years post-bariatric surgery. While Pcpe1 mRNA levels were either unchanged or increased.^67,68^ The statistically significant correlation between Pcpe2 mRNA and weight loss was consistent with earlier reports from our lab using an independent study of pre- and post-bariatric surgery patients.^50^ Furthermore, additional data from participants stratified into metabolically healthy lean (MHL), metabolically healthy obese (MHO), and metabolically unhealthy obese (MUO) **(Figures 1E-F and Supplemental Figure E)** were analyzed.^62^ Here, *PCOLCE2* mRNA levels were significantly higher in MUO than MHL and MHO groups, with no difference between the 2 metabolically healthy groups **(Figure 1E).** These data strongly suggest that metabolic health, rather than simply obesity, relates to the change in Pcpe2 mRNA abundance. Comparatively, *PCOLCE1* mRNA abundance did not differ significantly across groups, suggesting a different relationship between expression and adipose tissue mass **(Supplemental Figure 1E)**. Furthermore, a subset of 6 participants who had undergone Roux-en-Y gastric bypass surgery and 9 participants who consumed a low-calorie diet and lost ∼18% of body weight were analyzed and showed a statistically significant decrease in *PCOLCE2* mRNA abundance with weight loss **(Figure 1F),** which is consistent with an earlier published study^50^ suggesting that elevated *PCOLCE2* may serve as a biomarker of unhealthy adipose tissue expansion and remodeling.

Next, the effects of diet on the cellular expression of Pcpe2 mRNA were investigated. Data from a published study^25^ in which the distribution of adipocyte PCs and their gene expression were compared using two different diets. A UMAP from an sc-RNA-seq dataset shows mouse adipose PCs from epididymal (EAT) and inguinal (IAT) adipose tissue after a standard diet (SD), or 2 and 10 weeks of WD **(Figure 1H, left panel; Supplemental Figure 1H, left panel).** Here, both diet and duration of WD significantly altered cellular distribution. Specifically, a redistribution of *Pcolce2* mRNA expression (**Figure 1H, right panel)** compared to *Pcolce1* mRNA (**Supplemental Figure 1H, right panel)** across different dietary conditions reveals qualitatively distinct patterns. In EAT and IAT plots, *Pcolce2* mRNA shows higher expression in nearly all PC populations, with the highest at 10-week WD IAT, while *Pcolce1* mRNA shows a significantly different pattern of expression in both EAT and IAT. These data suggest that Pcpe2 expression increases in PC populations during WD feeding, whereas Pcpe1 expression may not, suggesting that Pcpe1 and Pcpe2 may play distinct, non-overlapping roles in obesity-associated adipose tissue expansion and remodeling.

### Adipose Tissue-Specific Pcpe2 Deletion Protects against WD-Induced Obesity

To experimentally investigate the role of Pcpe2 in adipose tissue expansion and remodeling, we generated adipose tissue-specific Pcpe2 knockout mice (TgAdip^+^Pcpe2^KO^) using the specific adiponectin Cre (Adip) and compared them to their adiponectin Cre-negative littermate controls (TgAdip^-^Pcpe2^Ctrl^). The terms VAT refer to mouse peri-gonadal fat; SAT to inguinal fat; and BAT to interscapular fat. During 20 weeks of consuming a WD, the body weights of female and male mice were monitored **(Figure 2A-E and Supplemental Figure 2A-B)**. At the end of the 20 weeks, images of the body and fat pads were taken, and the final total body and fat pad weights were measured. A reduction in TgAdip^+^Pcpe2^KO^ body weight relative to control mice was accompanied by reductions in VAT, SAT, and BAT pad weights (**Figure 2D-E, Supplemental Figure 2B).** Also consistent with the reduction in body and fat pad weight, TgAdip⁺Pcpe2^KO^ mice exhibited significantly improved glycemic control **(Figure 2F-G, Supplemental Figure 2C-D)**. Intraperitoneal glucose tolerance test (GTT) showed that both male and female TgAdip⁺Pcpe2^KO^ mice had improved glucose clearance, with the area under the curve (AUC) for glucose excursion significantly reduced in TgAdip⁺Pcpe2^KO^ mice relative to controls. Consistent with this, plasma insulin levels were also significantly lower in WD-fed TgAdip⁺Pcpe2^KO^ mice **(Figure 2H),** suggesting enhanced insulin sensitivity with the loss of Pcpe2 in adipose tissue.

**Figure 2.**
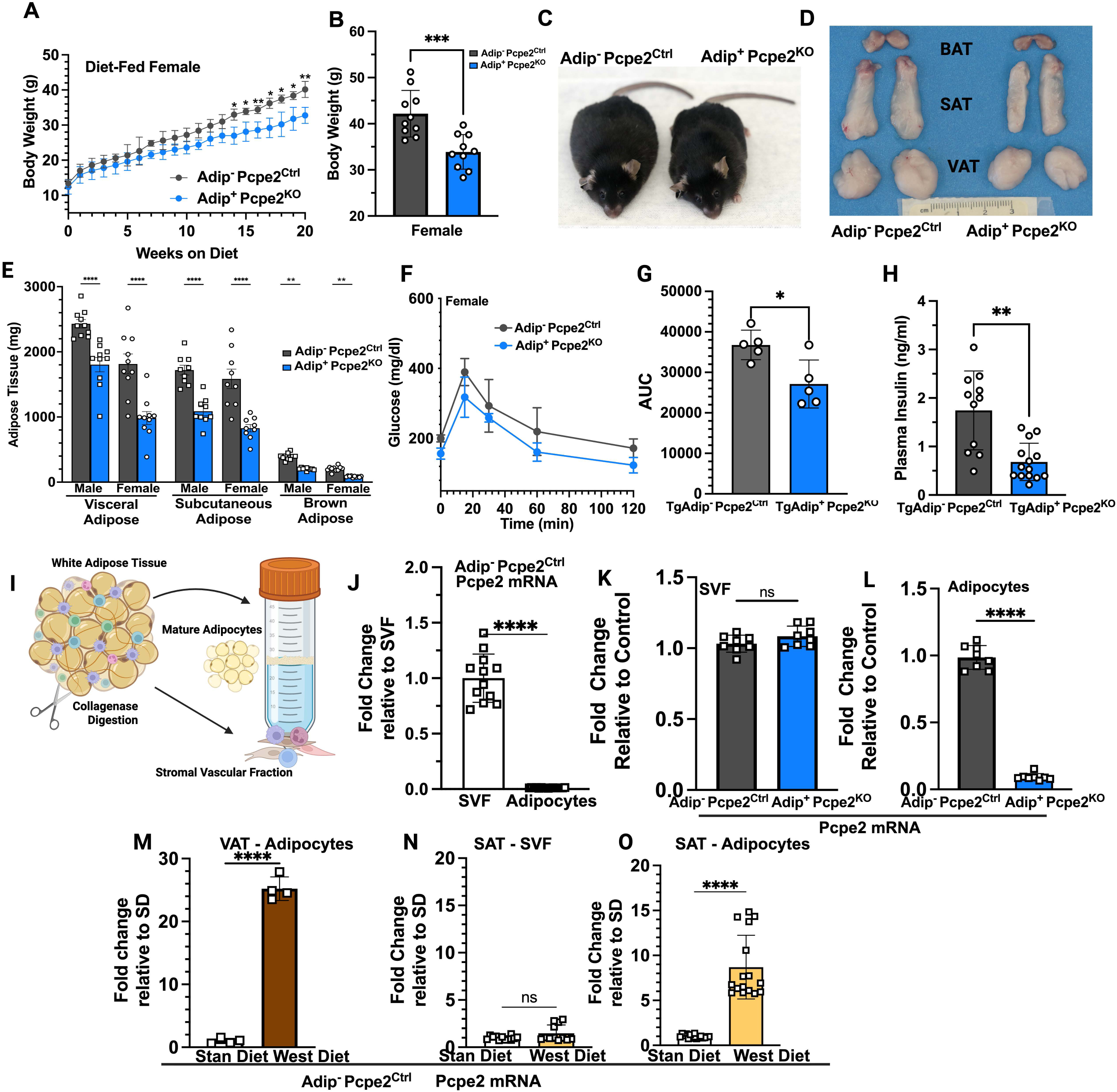
Adipose Tissue-Specific Pcpe2 Deficiency Protects Against Diet Induced Obesity. Adipose tissue-specific Pcpe2 knockout (Adip^+^Pcpe2^KO^) and floxed littermate controls (Adip Pcpe2^Ctrl^) mice were fed a WD for 20 weeks **(B)** End point body weight in female mice of indicated genotype **(C)** Representative images of Adip^+^Pcpe2^KO^ and Adip^-^ Pcpe2^Ctrl^ mice showing whole body appearance **(D)** Representative images of excised visceral (VAT), subcutaneous (SAT), and brown (BAT) adipose depots from indicated genotype **(E)** End point pad weights for VAT, SAT, and BAT from male and female mice of indicated genotype fed WD for 20 weeks **(F)** Glucose tolerance test (GTT) of WD female mice of indicated genotype **(G)** Area under the curve (AUC) for GTT **(H)** Fasted plasma insulin levels for WD-fed female mice **(I)** Illustration showing isolation of mature adipocytes from stromal vascular fraction (SVF) **(J)** Adip^-^Pcpe2^Ctr^ WAT Pcpe2 mRNA levels in SVF vs adipocyte fraction from standard diet (SD) fed mice **(K)** WAT SVF Pcpe2 mRNA from SD fed mice from indicated genotype **(L)** WAT adipocyte Pcpe2 mRNA levels from SD fed mice from indicated genotype **(M)** VAT adipocyte Pcpe2 mRNA levels comparing SD vs WD fed Adip^-^Pcpe2^Ctrl^ mice **(N)** SAT SVF Pcpe2 mRNA levels of SD vs WD fed Adip^-^Pcpe2^Ctrl^ mice **(O)** SAT adipocyte Pcpe2 mRNA levels of SD vs WD fed Adip^-^Pcpe2^Ctrl^ mice. All data shown are the mean ± SEM, from age and sex matched mice (n=11-15). Data were analyzed using unpaired t-test or one-way ANOVA followed by Tukey’s post hoc test. *p < 0.05 **p<0.005.

To verify the specificity of adiponectin Cre recombinase expression, we examined VAT, SAT, BAT, liver, muscle, and heart tissues for Cre-mediated excision. Specific probes were designed to detect cleavage within the floxed Pcpe2 DNA alleles **(Supplemental Figure 2E)**. Control littermates show no probe signal in any tissues, while TgAdip⁺Pcpe2^KO^ mice show cleavage in BAT, SAT, and VAT, but not in liver, muscle, or heart tissue, verifying the specificity of the adiponectin Cre recombinase. Given the specificity of adiponectin Cre to adipose tissue, we also examined the effects of deleting Pcpe2 using the platelet-derived growth factor receptor alpha Cre (Pdgfra) recombinase Cre. In this case, Pcpe2 would be deleted in al PCs, upstream of adiponectin expression.^69^ Despite this Cre targeting Pcpe2 expression early in the adipocyte lineage, both Cre-recombinase strategies produced largely overlapping phenotypes, as shown by a similar reduction in body weight and VAT pad weights after 20 weeks of WD **(Supplemental Figure 2F-G)**.

We next created and examined mice overexpressing Pcpe2 in adipose tissue. These mice were generated in a tissue-specific manner by crossing Pcpe2^Ox^ mice with Pdgfra Cre or adiponectin Cre to obtain PC^+^Pcpe2^Ox^ and their negative littermate controls. Examining Pcpe2 mRNA expression in VAT revealed approximately 8-fold higher levels in these mice **(Supplemental Figure 2H).** The expression specificity was also investigated, and excision was observed only in PC^+^Pcpe2^Ox^ mouse VAT, SAT, and BAT **(Supplemental Figure 2I)**. PC^+^Pcpe2^Ox^ mice and controls were fed WD for 20 weeks, after which body and pad weights were measured and compared to adipose-specific Pcpe2 deletion mice **(Supplemental Figure 2J-L).** Histological comparison of VAT from 20-week WD-fed control, PC⁺Pcpe2^Ox^, and PC⁺Pcpe2^KO^ mice **(Supplemental Figure 2L)** using Masson’s Trichrome staining was performed. Consistent with the results comparing body and fat pad mass, histology showed that hypertrophic adipocytes were attenuated when Pcpe2 was deleted, suggesting a reciprocal relationship between Pcpe2 expression and adipose expansion, consistent with data shown in **Figure 1D-F** and a previously published study.^50^

Next, the relative cellular contribution of Pcpe2 expression between WAT PCs in SVF and mature adipocytes was examined. WAT from SD-fed TgAdip^-^Pcpe2^Ctrl^ mice was digested with collagenase, and the SVF was separated from mature adipocytes using previously published methods and illustrated^55^ **(Figure 2I)**. RT-PCR was performed on extracted VAT **(Figure 2J)** and SAT RNA (data not shown), and showed that in VAT and SAT, the SVF expresses significantly more Pcpe2 mRNA than mature adipocytes. Next, we compared the expression of Pcpe2 mRNA in SVF between SD-fed TgAdip^-^Pcpe2^Ctrl^ and TgAdip⁺Pcpe2^KO^ mice using age- and sex-matched tissue **(Figure 2K).** Using either WAT or SAT no significant difference was noted in Pcpe2 mRNA expression in SVF. However, as expected, when Pcpe2 mRNA abundance was compared in mature adipocytes between SD-fed TgAdip^-^Pcpe2^Ctrl^ and TgAdip⁺Pcpe2^KO^ mice, a significant reduction in Pcpe2 mRNA expression was noted in Tg⁺Pcpe2^KO^ mice **(Figure 2L),** confirming genotyping results and the specificity of the adiponectin Cre.

To examine how WD affects Pcpe2 mRNA expression in WAT, RNA from mature adipocytes derived from TgAdip^-^Pcpe2^Ctrl^ mice fed either SD or WD was compared **(Figure 2M)**. Pcpe2 mRNA was elevated ∼25-fold in VAT adipocytes in response to WD. Next, Pcpe2 mRNA in SVF and mature adipocytes was compared for their response to WD using TgAdip^-^Pcpe2^Ctrl^ mice fed either SD or WD **(Figure 2N-O).** In SAT-derived SVF, no change in Pcpe2 mRNA abundance was noted between SD and WD **(Figure 2N)**. On the other hand, an ∼8-fold increase in SAT adipocytes expressing Pcpe2 mRNA was noted when SD levels were compared to WD levels (**Figure 2O)**. Thus, both mature adipocytes from VAT and SAT, show a remarkable upregulation of Pcpe2 mRNA expression in response to WD. These findings align well with observations of changes in Pcpe2 mRNA abundance in patients undergoing weight gain and loss. Taken together, these data suggest that Pcpe2 expression in WAT adipocytes is highly responsive to WD and likely contributes to the unhealthy expansion and remodeling of adipose tissue.

### WAT Beiging and Energy Expenditure in Adipose Tissue-Specific Pcpe2 Knockout Mice

Studies were conducted to investigate how deleting Pcpe2 in adipose tissue protects against WD-induced obesity. Histological analyses of VAT from 20-week SD- and WD-fed female TgAdip^-^Pcpe2^Ctrl^ (**Figure 3A, top panels**) and TgAdip⁺Pcpe2^KO^ (**Figure 3A, bottom panels**) mice were conducted. VAT pad sections were stained with Masson’s Trichrome, and images (20X) revealed that for both genotypes VAT adipocytes enlarged from SD to WD-fed mice **(Figure 3A left vs right panels).** But strikingly, TgAdip⁺Pcpe2^KO^ mice displayed generally smaller adipocytes with regions of smaller adipocytes consistent with WAT beiging **(Figure 3A, bottom left and right).** VAT sections were also imaged at 2X magnification (**Supplemental Figure 3A),** showing numerous foci containing smaller beige adipocytes within the VAT pad. While VAT from WD-fed TgAdip^-^Pcpe2^KO^ mice showed regions of beige adipocytes, the BAT depot showed evidence of “whitening” of brown adipocytes, consistent with lipid accumulation **(Supplemental Figure 3B).** After 20 weeks of WD, both TgAdip^-^Pcpe2^Ctrl^ and TgAdip⁺Pcpe2^KO^ BAT sections were stained with Masson’s Trichrome and showed a noticeable difference in lipid accumulation, compared with sections from 20-week SD-fed mice (**Supplemental Figure 3B, left top and bottom panel).** Furthermore, WD-fed TgAdip^-^Pcpe2^Ctrl^ mouse BAT appeared to have a greater extent of whitening when compared to TgAdip⁺Pcpe2^KO^ mouse BAT sections (**Supplemental Figure 3B, right top and bottom panels).**

**Figure 3.**
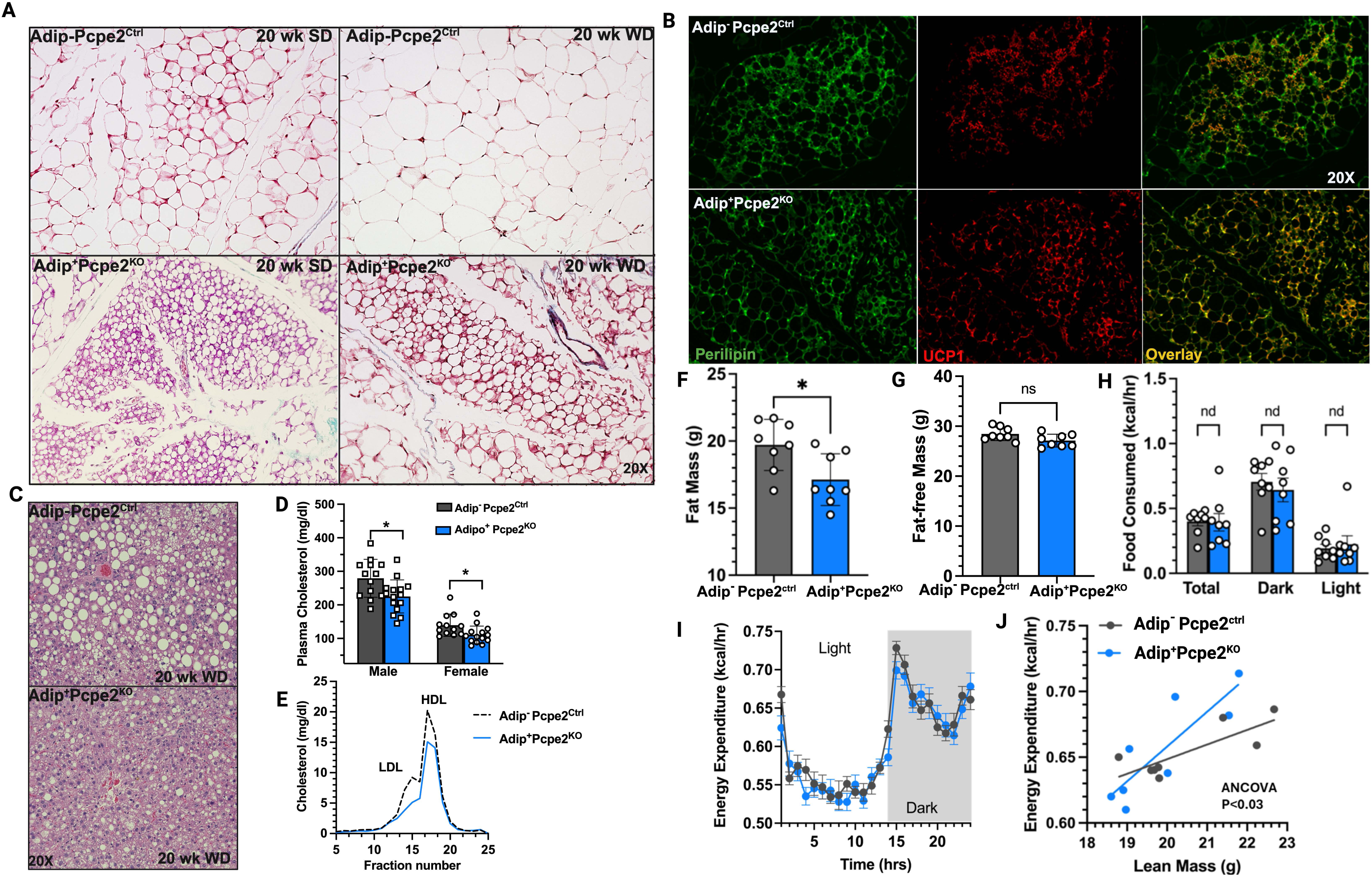
Pcpe2 Deficiency Enhances Energy Expenditure and WAT Beiging. **{A}** Representative Masson’s Trichrome stained VAT images from female Adip^-^Pcpe2^Ctrl^ **(top panels}** and Adip^+^Pcpe2^KO^ **{bottom panels}** mice fed SD **{left panels}** or WO **{right panels}** for 20 weeks **{B}** Representative immunofluorescence images of VAT from 20 week WO-fed Adip^-^Pcpe2^Ctrl^**(top panels}** and Adip^+^Pcpe2^KO^ **{bottom panels}** mice **{left panels}** immunostained for lipid droplet-perilipin (green), and **{center panels}** UCP1 (red), with **{right panels}** merged colocalization. **{C}** Representative images of 20 week WO-fed liver sections stained with H&E from indicated genotypes **{D}** Plasma cholesterol concentrations from male an female mice of indicated genotype fed WO **{E}** FPLC separation of plasma from WO-fed mice showing lipoprotein cholesterol concentration per fraction for indicated genotypes **{F-G}** Fat and fat-free mass from indicated genotype, determined by EchoMRI analysis from 20 week female WO-fed mice **{H}** Food intake during light and dark cycles using Promethion metabolic cages {I} Energy expenditure during light and dark cycle **{J}** Energy expenditure anaylzed by CalR using GLM and ANCOVA. Data represent mean± SEM (n=8-15) Statistical significance was determined by unpaired t-test or ANOVA.

To examine beiging, VAT sections from WD-fed TgAdip^-^Pcpe2^Ctrl^ and TgAdip⁺Pcpe2^KO^ mice were stained for UCP1 **(Figure 3B)** and counterstained with perilipin for immunofluorescence imaging. TgAdip^-^Pcpe2^Ctrl^ mice (**Figure 3B top**) show relatively less UCP1 staining compared to WD-fed TgAdip⁺Pcpe2^KO^ **(Figure 3B bottom)**. Higher levels of UCP1 expression in TgAdip⁺Pcpe2^KO^ mice were also supported by RT PCR analysis to determine the level of UCP1 mRNA (**Supplemental Figure 3C).** A 3-4-fold increase in UCP1 mRNA abundance was observed in TgAdip^-^Pcpe2^KO^ SAT when compared to control mice. Western blot analysis confirmed this finding, as UCP1 expression in SAT protein extracts showed ∼2-fold increase in intensity in Adip⁺Pcpe2^KO^ mice **(Supplemental Figure 3D-E).** Also, Western blot analysis of Pparg levels, the master regulator of adipogenesis, was carried out **(Supplemental Figure 3F-G**) and showed that TgAdip⁺Pcpe2^KO^ SAT has ∼1.3-fold increase in Pparg levels compared to control mice. Therefore, taken together, histology, mRNA abundance and UCP1 protein abundance strongly support the idea that WAT beiging is upregulated in TgAdip⁺Pcpe2^KO^ mice compared to controls.

Next, liver sections from 20-week WD-fed TgAdip^-^Pcpe2^Ctrl^ and TgAdip⁺Pcpe2^KO^ **(Figure 3C)** mice were compared. Sections were stained with H&E and showed a markedly different degree of lipid accumulation, with TgAdip⁺Pcpe2^KO^ showing fewer and smaller lipid droplets compared to controls. Interestingly, consistent with a decreased liver lipid accumulation, the plasma cholesterol concentration was also lower in WD-fed male and female Adip⁺Pcpe2^KO^ mouse plasma **(Figure 3D)**. Additionally, FPLC fractionation of plasma to separate low-density lipoprotein (LDL) and high-density lipoprotein (HDL) showed that TgAdip⁺Pcpe2^KO^ mice have lower levels of both LDL and HDL cholesterol **(Figure 3E)** in plasma. These findings raise the question of how Pcpe2 deletion in adipocytes affects atherosclerosis progression, since published studies have shown that thermogenic stimulation promotes HDL-cholesterol clearance and increases macrophage-to-feces reverse cholesterol transport in mice.^70^ Other studies have shown that the VLDL-VLDL receptor axis in brown adipocytes plays a major role in regulating thermogenesis and plasma VLDL/LDL concentrations.^71^

We next conducted metabolic phenotyping on 20-week WD-fed female TgAdip-Pcpe2^Ctrl^ and TgAdip⁺Pcpe2^KO^ mice using the Promethion metabolic cage system (**Figure 3F-J)**. TgAdip⁺Pcpe2^KO^ mice exhibited significantly lower body weight, consistent with previous measures, and showed no significant difference in fat-free mass **(Figure 3F-G)** or lean mass (data not shown). No significant difference in food intake was seen between genotypes **(Figure 3H)**. As expected, the lower body weight in TgAdip⁺Pcpe2^KO^ mice affected these measurements; however, after a generalized linear model was generated using the CalR analytical portal^58^, Pcpe2-deficient mice showed a statistically significant elevation in energy expenditure compared to control mice. Thus, collectively these data are consistent with the concept that deletion of Pcpe2 in WAT increases beiging.

### Pcpe2, WAT Inflammation and Fibroadipogenic Progenitors (FAPs)

In response to dietary caloric overload, WAT expansion can be limited, and inflammation ensues. To investigate the role Pcpe2 plays in this process, immunofluorescence of CD68^+^ immune cells in VAT sections from WD-fed TgAdip-Pcpe2^Ctrl^ and TgAdip⁺Pcpe2^KO^ mouse tissue was conducted **(Figure 4A)**. Control mice exhibited prominent CD68⁺ macrophage infiltration surrounding enlarged adipocytes **(Figure 4A, left),** shown at 10X, and 40X magnification, while WD-fed TgAdip⁺Pcpe2^KO^ mice showed reduced CD68^+^ intensity **(Figure 4A, right).** RT PCR analysis was conducted on RNA derived from WD-fed TgAdip-Pcpe2^Ctrl^ and TgAdip⁺Pcpe2^KO^ VAT tissue **(Figure 4B),** including Il6, Ccl2, Hif1a, Tnfa, Fn1, Tgfb, and Col1 genes. The expression of these inflammatory VAT markers was significantly reduced compared to controls. These findings raise interesting questions, given that adipose tissue contains several populations of adipocyte PCs with distinct inflammatory properties.^18,21,72^ Thus, to further investigate, VAT SVF progenitor populations were quantified by FACS analyses following collagenase digestion and separation of SVF from mature adipocytes **(Figure 4C**). After exclusion of CD45⁺ immune cells and CD31⁺ endothelial cells, APCs were identified as CD140b⁺ (Pdgfrβ⁺) and further resolved based on expression of the surface markers Ly6C and CD9 **(Figure 4D).**^18,20^ FACS profiling demonstrated a pronounced shift in progenitor composition in VAT comparing TgAdip⁺Pcpe2^KO^ mice with controls. Pcpe2-deficient VAT PCs displayed a significant reduction in the proportion of inflammatory FAPs, accompanied by a relative enrichment of APCs, indicative of a less inflammatory and more adipogenic niche **(Figure 4E-F)**. Quantitative analysis revealed that SD-fed TgAdip⁺Pcpe2^KO^ mice exhibited a higher percentage of APCs with a corresponding reduction in FAPs compared with controls **(Figure 4E).** This shift was further accentuated after WD-feeding, in which control mice showed a robust expansion in the percentage of FAPs, while Pcpe2-deficient mice had a more modest shift **(Figure 4F).**

**Figure 4.**
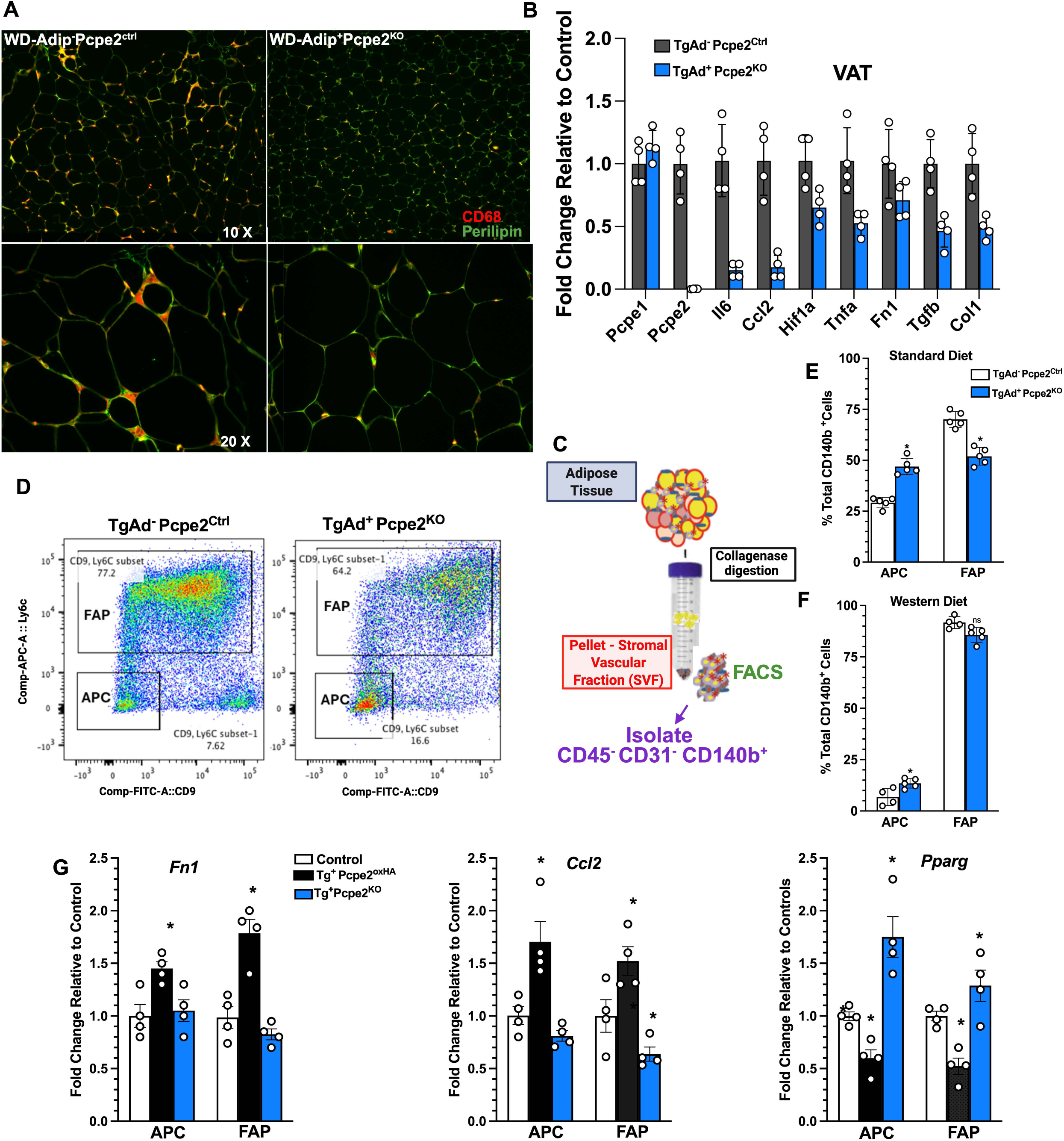
Loss of Pcpe2 in WAT Attenuates Distribution of Fibroinflammatory Adipocyte Precursors and Inflammation. Representative WAT images stained for CD68 and Perilipin from mice fed WO for 20 weeks **(A, left panels)** Adipo^-^Pcpe2^Ctrl^ and **(A, right panels)** Adipo^+^Pcpe2^KO^ mice, at **(A, top panels),** 1OX, and **(A, bottom panels)** 40X magnifications **(B)** WO-fed VAT mRNA abundance for selected inflammatory genes following RT PCR **(C)** Isolation of SVF for FACS analyses **(D)** Representative FACS dot plot showing Ly6c and CD9 gates defining FAPs and APCs following gating for co45+ and CD31+ and CD140b+ (Pdgfrb+) **(E-F)** Percent of CD140b+ cells for **(left)** FAPs **(right)** APCs shows change in percentage comparing SD to WO-fed mouse VAT and relative to genotype **(G)** WO VAT derived APC and FAP were isolated and their RNA analyzed by RT PCR for selected genes for mRNA abundance. Data shown represent the mean ± SD, n = 4-6 mice per genotype. Statistical analyses were performed using ANOVA with Tukey’s multiple comparisons and unpaired t-tests.

Next, APCs and FAPs from WD-fed TgAdip^-^Pcpe2^Ctrl^, TgAd^+^Pcpe2^Ox,^ and TgAdip⁺Pcpe2^KO^ were isolated, and RNA was analyzed by RT-PCR. Here, the expression of pro-fibrogenic and inflammatory genes **(Figure 4G)** was investigated. APCs and FAPs from Pcpe2-overexpressing (TgAd^+^Pcpe2^Ox^) mice displayed significantly increased expression of key inflammatory and fibrogenic mediators, including Fn1, Ccl2. While Pcpe2-deficient APCs and FAPs showed a significant increase in the expression of the adipogenic mediator, Pparg, compared with control mice. It is known that FAPs express ∼2-fold higher levels of TGFbR1 and 2 receptors than other PCs in adipose, likely contributing to adipose tissue containing the highest levels of TGFBR2 among all tissues.^66^ TGFB1 is also associated with obesity and insulin resistance in both animals and humans.^73^ Both Insulin and high glucose are known to stimulate TGFb1 receptor to the cell surface.^74^ Sorted VAT FAPs were cultured *ex vivo* and treated with 2 ng/ml of TGFb for 3 hours. Conditioned medium was collected and probed for TGFb1 using Western blot analysis and showed that FAPs TgAd^+^Pcpe2^Ox^ cells responded with a significantly higher accumulation of TGFb1 ligand **(Supplemental Figure 4A)** over controls. Overexpression of Pcpe2 and its association with markers of inflammation were also supported by studies in which VAT pads from TgAdip^-^Pcpe2^Ctr^ and TgAd^+^Pcpe2^Ox^ mice were transplanted into Adiponectin^Cre+^ DTA^fl/+^ mice.^59^ These lipoatrophic mice lack WAT from birth and have only bone marrow adipocytes, exhibiting dysregulated glucose tolerance and elevated plasma triglyceride levels. Following VAT transplantation, no significant differences between groups were noted in body weight, while serum triglycerides normalized within a few weeks. After ∼10 weeks, the transplanted fat VAT pads were removed and analyzed by histology **(Supplemental Figure 4C).** Sections were first analyzed for the presence of the Pcpe2 HA-tag, confirming Pcpe2 overexpression in adipose. Additional sections were examined by staining with Masson’s trichrome and CD68/perilipin **(Supplemental Figure 4C)**. These data confirm the cell-intrinsic nature of Pcpe2 overexpression on inflammation, given the higher CD68 signal compared to controls. These findings strongly suggest that Pcpe2 not only regulates PC cell composition but also governs the inflammatory and fibrogenic state of PCs in VAT, likely through local signaling pathways involving TGFb.

### Mitochondrial Function and Ex Vivo Adipocyte Differentiation

To investigate how Pcpe2 expression shifts the balance of WAT beiging towards inflammation and fibrosis, studies were conducted using *ex vivo* differentiated adipocytes from mouse adipose tissue. A timeline showing the steps involved in differentiating SVF-derived PCs is shown (**Figure 5A).** PCs from SD-fed SAT are cultured and, after 4 days of differentiation to allow adipogenesis/differentiation, wells of cells were stained with the lipid stain, Bodipy, and representative images were captured **(Figure 5B)**. In this context, the greater extent of lipid staining **(Figure 5 B right panels)** indicates a higher degree of adipogenesis in cells lacking Pcpe2. Thus, a statistically significant adipogenic potential (area of lipid staining per cell) from Pcpe2^KO^ adipocytes relative to control cells **(Figure 5C)** reflects a greater degree of differentiation. Another contributor to adipogenic potential is lipid uptake. We examined the ability of differentiated adipocytes to take up DiI-VLDL **(Figure 5D)** and found that, in the absence of Pcpe2, uptake was greater and accompanied by increased VLDL receptor expression **(Figure 5E)** compared to control adipocytes.

**Figure 5.**
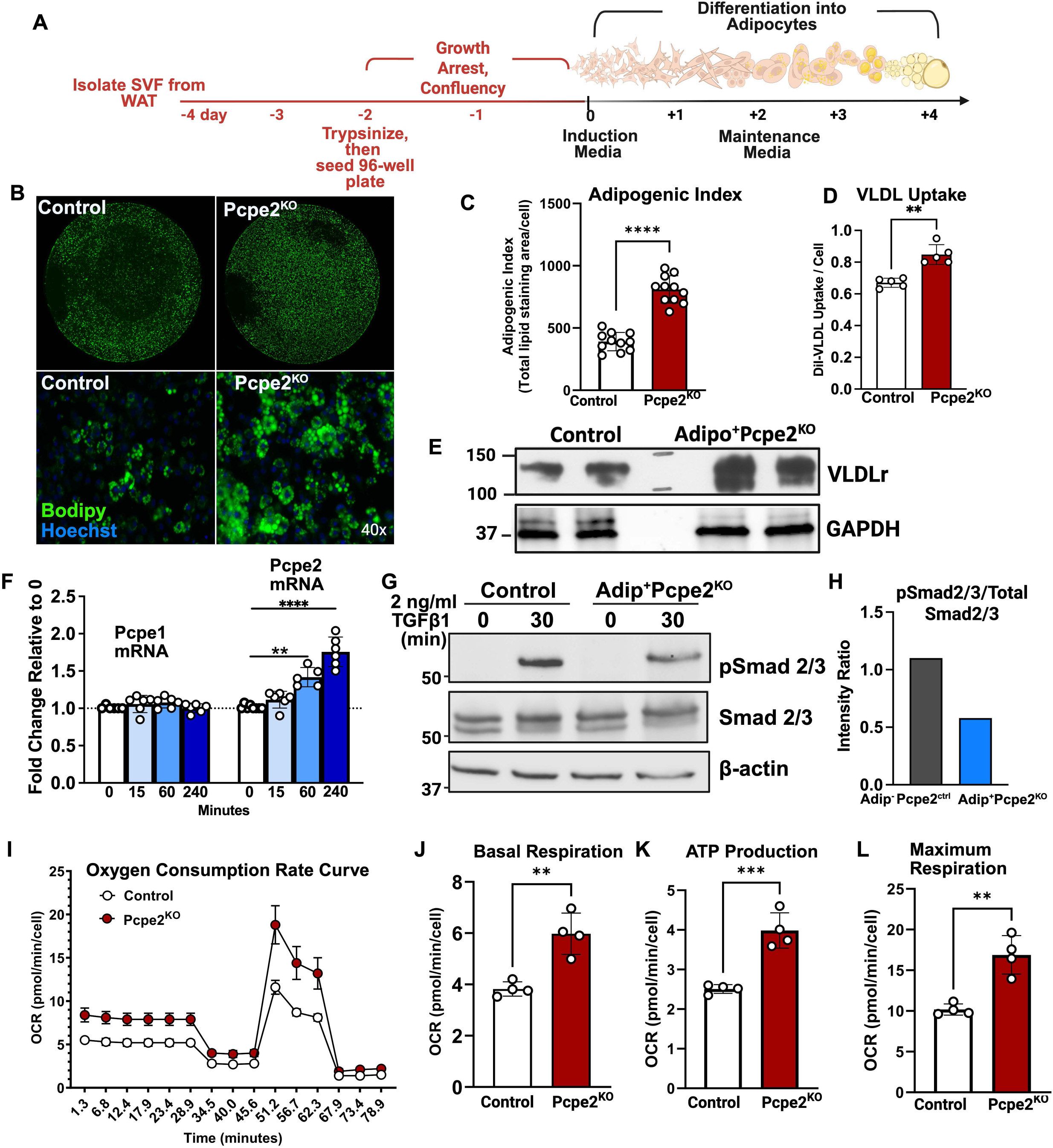
Pcpe2 Expression Regulates Adipocyte Differentiation and Mitochondrial Efficiency. **(A)** Illustration depicting timeline of differentiating SVF cells into adipocytes *ex vivo* **(B)** Representative images of differentiated adipocytes stained with Bodipy (lipid) and Hoechst (nuclear) **(top panels)** shows single 96 well from rom control and Pcpe2^KO^ cell **(bottom panels)** image taken from well at 40X magnification **(C)** Adipogenic index of differentiated adipocytes measures total Bodipy lipid stained area per cell **(D)** Uptake of Dil-VLDL by differentiated adipocytes for indicated genotype **(E)** Western blot for VLDL receptor abundance in differentiated adipocyte protein extracts **(F)** TGFb treatment (2 ng/ml) of differentiated adipocyte RNA from control mice following RT PCR **(G)** Western blot for phospho-Smad2/, total Smad2/3 and b-actin from differentiated adipocytes treated for 30 min with 2 ng/ml of TGFb **(H)** Quantification of band intensity for pSmad2/3 to total Smad2/3. (I) Seahorse study from differentiated adipocytes showing oxygen consumption rate (OCR) **(J)** Basal Respiration **(K)** ATP Production **(L)** Maximum Respiration. All data represent mean ± SEM of differentiated adipocytes from SAT SVF with n=6 wells per genotype -96 well plate. Statistical significance was determined by unpaired t-test or ANOVA.

Pcpe2 has been described as an ECM resident glycoprotein, containing more O-linked than N-linked glycosylation.^35^ A closer examination of Pcpe2’s amino acid sequence^32,65^ identifies a linker region with 9 threonines, highly conserved, which are potential sites for O-linked glycosylation. Studies were performed to more finely identify the glycans and their potential function. To do this, we used our previously described mammalian expression system^50^ to express both the wild-type PCPE2 and a mutant of wild-type PCPE2, abbreviated as NG PCPE2. This clone contains substitutions of 9 linker-region threonine’s with alanine’s to effectively block O-linked glycosylation at those sites. To confirm this, wild-type and NG PCPE2 glycoproteins were analyzed by SDS-PAGE and stained with colloidal blue **(Supplemental Figure 5A).** NG-PCPE2 carries threonine-to-alanine substitutions at the nine O-glycosylation sites of the mucin-like linker domain introduced above, removing O-glycosylation at those positions. To confirm this, wild-type and NG PCPE2 glycoprotein were analyzed by SDS-PAGE and stained with colloidal blue **(Supplemental Figure 5A).** Mass spectrometry confirmed O-glycan occupancy at the nine-linker threonine’s of wild-type PCPE2, resolved the dominant glycan structures detected at each occupied site, and confirmed loss of occupancy across all nine sites in NG-PCPE2 (**Supplemental Table 1, Supplemental Figure 5B**). To our knowledge, this is the first site-resolved map of PCPE2 O-glycosylation To determine the effect of removing PCPE2 linker glycosylation, SVF PCs from Pcpe2^KO^ mice was differentiated, with a final concentration of 150 ng/ul of NG PCPE2 added to the culture medium at each media change, beginning at induction and continuing through the final 4 days of differentiation. On the final day of differentiation, cells were analyzed for their adipogenic index (**Supplemental Figure 5C).** In adipocytes that received no additions, as seen previously **(Figure 5C)**, SVF cells differentiated to a greater extent in the absence of Pcpe2. However, when NG PCPE2 was added, the adipogenic index was significantly reduced and was near control values. DiI-VLDL uptake was also tested in differentiated adipocytes (**Supplemental Figure 5D).** In these experiments, the addition of PCPE2 at 150 ng/ul reduced VLDL uptake, while the addition of NG PCPE2 showed some attenuation in uptake that was not statistically significant. Glycosylated PCPE2, but not NG-PCPE2, significantly reduced DiI-VLDL uptake, indicating that the effect of PCPE2 on adipocyte lipid uptake depends on O-glycosylation of the mucin-like linker domain. Taken with the differentiation data, these results indicate that the linker-domain O-glycans are required for at least part of PCPE2 activity in adipocytes and support a glycan-dependent role for Pcpe2 in adipose remodeling.

Previous studies have suggested that a relationship between WAT fibrosis and beiging is heavily influenced by TGF-β signaling^75^ and has been extensively studied.^22,23,76^ One study exploring this connection found that Pcpe2 mRNA expression, but not Pcpe1 mRNA, in mouse BAT was highly responsive to c treatment.^77^ Other reports show that TGF-β signaling suppresses beige adipogenesis by recruiting dedicated beige progenitors and regulating the balance between adipose tissue fibrosis and beige adipogenesis.^75^ To study this relationship, we treated day 4 differentiated adipocytes with 2 ng/ml TGF-β1 for various times **(Figure 5F)**. RNA was isolated and amplified by RT PCR to determine the abundance of Pcpe1 and 2 mRNA in response to treatment. By 60 min, Pcpe2 mRNA was significantly increased, and by 240 min, there was ∼2-fold increase in mRNA expression, with no change in Pcpe1 mRNA abundance. Since TGF-β1 signals through the pSmad 2/3^73^ pathway, Western blot analysis of differentiated adipocyte extracts was probed for pSmad 2/3 **(Figure 5G-H).** Adipocytes that lacked Pcpe2 showed a significant attenuation of pSmad 2/3 relative to total Smad after 30 min of TGF-β1 treatment, suggesting that the ECM glycoprotein may participate in TGF-β1 signaling, possibly by altering TGFBR1 and 2 interaction or availability.^78^

Next, differentiated adipocytes from control and Pcpe2^KO^ mouse SAT were studied to determine their mitochondrial function using Seahorse extracellular flux analyses **(Figure 5I-L)**. Pcpe2-deficient adipocytes displayed higher oxygen consumption rates (OCR) during the mitochondrial stress test, indicating elevated mitochondrial respiration **(Figure 5I)**. Furthermore, quantitative analyses revealed a significant increase in basal respiration, ATP production, and maximal respiration **(Figure 5J-L).** Additionally, spare respiratory capacity and proton leak were also increased (data not shown). Glycolytic function was also measured **(Supplemental Figure 5E-H)** and indicated that Pcpe2 expression did not have a significant difference on glycolytic function in differentiated adipocytes as measured by the extracellular acidification rate (ECAR). Additionally, Undifferentiated SVF cells were also studied and compared to the *ex vivo* differentiated cells using Seahorse extracellular flux analyses **(Supplemental Figure 5I-L).** In cells lacking Pcpe2, both basal respiration and ATP production showed significant elevation over controls, but overall, at a much-reduced level compared to values obtained from adipocytes. Overall, these results strongly suggest that mitochondrial function is enhanced at the cellular level in the absence of Pcpe2.

## Discussion

Adipose tissue ECM O-glycoprotein, Pcpe2 has many distinct structural and functional characteristics that set it apart from its paralog, Pcpe1.^31,32,65,79^ In this report, we specifically demonstrate new and important roles for Pcpe2 during adipose tissue expansion and remodeling. Pcpe2, expressed in mature adipocytes, increases in response to WD and TGFb, both contributing to inflammation and fibrosis in human and mouse obesity. Our studies clearly show that adipose-specific, TgAd^+^Pcpe2^KO^ mice are resistant to WD-induced obesity and exhibit reductions in body and fat pad mass. Promethion metabolic cage studies demonstrated no significant differences in lean mass or food intake, yet WD-fed TgAd^+^Pcpe2^KO^ mice exhibited significantly higher energy expenditure than negative Cre littermate controls. WD-fed TgAd^+^Pcpe2^KO^ mice also showed reduced plasma glucose and lipoprotein cholesterol concentrations as well as reduced liver lipid content compared to controls, demonstrating an improved cardiometabolic profile. Markers of adipose tissue inflammation, such as Ccl2, Il6, and Tnfa mRNA, were also reduced in WD-fed TgAd^+^Pcpe2^KO^ adipose, while examination of VAT-derived CD140b^+^ PCs displayed a shift from FAPs, towards APCs, contributing to attenuation of local inflammation and fibrosis. In global Pcpe2^KO^ mice, a similar protection against WD-induced body and fat pad weight gain was seen.^50^ In the same report, *Pcolce2* mRNA abundance correlated with body fat in a dataset of obese human subjects undergoing bariatric surgery.^50^ Patients showed positive correlations for presurgery SAT *Pcolce2* mRNA abundance versus both body mass index and percent fat mass. After bariatric surgery-induced weight loss, correlations between *Pcolce2* mRNA expression and measures of body fat persisted, suggesting a role for Pcpe2 in adipose expansion and storage in humans.^50^ In the current report, we again show that weight gain and loss across multiple human cohorts affect *Pcolce2* mRNA abundance and strongly indicate that this O-linked glycoprotein could serve as an important biomarker for unhealthy adipose remodeling.

Mechanism(s) explaining Pcpe2’s action were explored using *ex vivo* differentiation of WAT-derived PCs. Here, the loss of Pcpe2 enhanced adipocyte differentiation as measured by adipogenic index, as well as VLDL uptake via the VLDL receptor. Differentiated adipocytes also showed reduced TGFβ-like signaling via pSmad2/3 to total Smad2/3 ratio. Our study and others^77,78^ have shown that Pcpe2 is induced in response to TGFb induction. Overall, these data are consistent with studies showing that a high-fat diet activates the TGFb/Smad3 signaling pathway, promotes adipose fibrosis, and blocks WAT beiging.^75^ It has been shown previously that TGF-β signaling inhibits the differentiation of WAT PCs via pSmad2/3 signaling and drives fibrosis, inflammation, and mitochondrial dysfunction, leading to insulin resistance^73,80^. Knockout of TGFBR1 promoted beige adipogenesis and protected against high-fat diet-induced obesity^81^. However, the biological actions of TFGb seem likely to vary with the local environment. In one study, TGFb appeared to be associated with insulin action and a Smad3-mediated positive feedback loop. These two mechanisms were likely coupled to the extracellular environment and to intracellular processes that stimulate the lipogenic pathway in adipocytes ^78^. These studies suggest that TGFb signaling in the ECM synchronizes intracellular lipid homeostasis to adapt to feeding-related insulin stimulation. In essence, showing that the signaling in the ECM can regulate intracellular lipogenic pathways, including FAK-AKT. Furthermore, our studies show elevated mitochondrial function, as measured by Seahorse assay on differentiated adipocytes in the absence of Pcpe2. These data are consistent with the idea that removing Pcpe2 from its role in enhancing TGFb signaling will lead to more WAT beiging. However, the precise mechanism by which Pcpe2 regulates the processes is currently under investigation.

Our studies also show, for the first time, that despite substantial structural homology^32^, Pcpe1 and 2 have markedly different O-glycosylation in their linker domains.^32^ While it was initially appreciated that these two glycoproteins had different glycosylation^35^, the current studies confirm the sites and shed light on the functionality of this linker region. Because O-glycans contained in a mucin-like domain commonly influence protein stability,^82^ localization, and binding interactions, the O-glycan-containing linker domain may provide a plausible structural basis for the functional divergence between the two paralogs. Typically, heavily O-linked domains like this one influence protein stability, localization, and binding interactions, thereby providing a structural basis for functional divergence. It is possible that this unique structural feature of Pcpe2 may explain why its adipose functions differently from those of Pcpe1. Pcpe2 carries a mucin-like linker domain with nine O-glycosylation sites that Pcpe1 lacks, as verified by mass spectrometry analysis. When all sites for O-glycation were removed, the protein NG-PCPE2, showed reduced adipogenic index and VLDL uptake compared to PCPE2. These observations indicate that the adipose activity of Pcpe2 is associated with O-glycosylation of the mucin-like linker domain rather than with the CUB or NTR domains shared between the paralogs, and they offer a structural explanation for the distinct, and in some cases opposing, behaviors of Pcpe1 and Pcpe2 noted throughout this study. They also raise the question of how these O-glycans act, whether by stabilizing the protein, directing its localization in the ECM, or modulating its interaction with TGFβ signaling components, which is a direction for future work.

Although evidence presented in this report focuses on the differences between Pcpe1 and Pcpe2, there are published studies suggesting that Pcpe1 also plays a role in promoting fibrosis, particularly in the heart and liver. In one study using global *Pcolce1*^KO,^ a significantly reduced liver fibrosis, body weight, and collagen I levels were noted, but no change in nonalcoholic steatohepatitis.^37^ This same group showed a significantly lower ejection fraction in *Pcolce1*^KO^ mice compared to controls but did not see any changes in cardiac collagen content or wound healing^38^. In other mouse models utilizing global and tissue-specific Pcpe1 knockout mice, the authors reported that Pcpe1 expressed in BAT circulates in plasma and promotes liver fibrosis and nonalcoholic steatohepatitis^39^. This study examined both global and BAT–specific deletions of Pcpe1 and found a marked reduction in liver fibrosis and circulating plasma Pcpe1 levels. Conversely, overexpression of Pcpe1 in brown adipose tissue exacerbated liver fibrosis, suggesting that high-calorie diet-induced stress increased Pcpe1 production BAT, thereby increasing circulating Pcpe1 levels and promoting liver fibrosis. In another report, this same group studied age-associated heart failure using global and BAT-specific *Pcolce1*^KO^ mice^83^. They concluded that Pcpe1 is a secreted extracellular matrix protein that acts as a BATokine, promoting fibrosis in the liver and heart during aging and obesity.

It seems clear from our work and that of others that more work is needed to fully appreciate the similarities and differences between these two very interesting glycoproteins. Collectively, our results show that the adipose tissue ECM O-glycoprotein Pcpe2 is a robust marker for unhealthy adipose tissue expansion and remodeling in humans and in mice. The expression of this glycoprotein contributes to inflammation and fibrosis associated with WD-induced obesity. Importantly, our data suggest that Pcpe2 expression inhibits white-to-beige adipogenesis by enhancing TGFβ-mediated pSmad2/3 signaling, thereby upregulating fibrosis. Continued studies to understand the mechanistic details of Pcpe2’s role in adipose expansion are currently underway.

## Acknowledgments

We appreciate the valuable assistance of Marjorie A. Kipp with technical support for our mouse colony and its maintenance. This research was supported by NIH RO1DK139164 (MST), RO1DK143978 (RKG), R01HL164460 (YC), R01DK101578, P30DK056341 (Nutrition Obesity Research Center), and UL1TR002345 (Washington University Institute of Clinical and Translational Sciences) and support from the Foundation for Barnes-Jewish Hospital (SK, GIS), RS-2024-00426031 (JKK), and F30CA281124 (RLM), and by an American Heart Association Postdoctoral Fellowship 26POST1557985 (MAB).

---

**Supplementary Figure 1.**
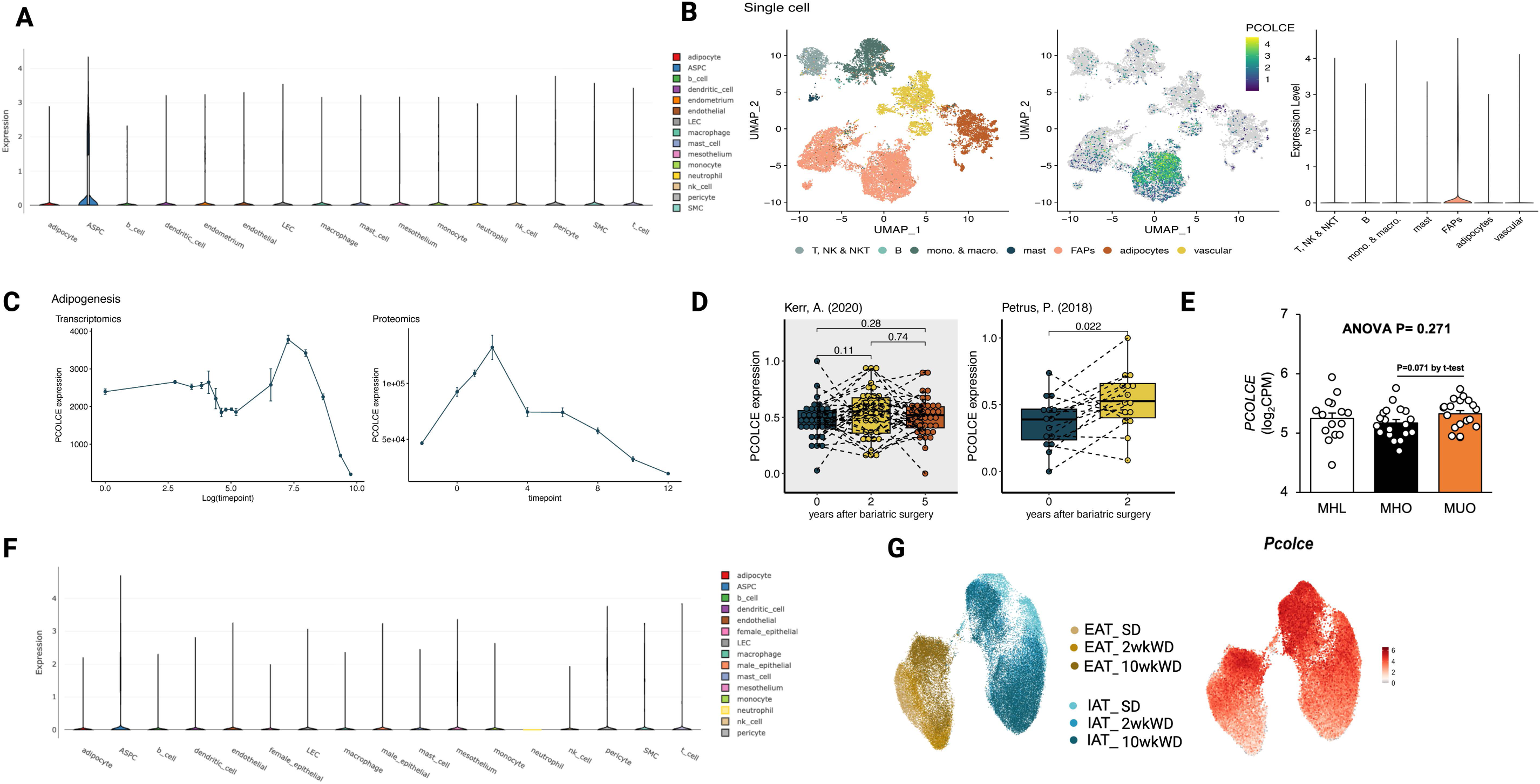
Expression of *PCOLCE I Pcolce* in Adipose Tissue in Humans and Mice. **(A)** *PCOLCE* relative expression among various cell types as determined by single-cell RNA sequencing (scRNA-seq) of human white adipose tissue (WAT) **(B)** UMAP of human WAT *PCOLCE* expression in fibroinflammatory adipocyte precursors (FAPs) and mature adipocytes relative to various immune cells **(C) (Left panel)** *PCOLCE* mRNA expression and **(Right panel)** PCPE protein expression expressed during adipocyte differentiation **(D)** Human *PCOLCE* mRNA expression from subcutaneous adipose tissue (SAT) collected before and after weight loss. Studies involved obese individuals who underwent bariatric surgery, with follow-up assessments conducted **(Left panel)** two and five years post-surgery as reported by Kerr et al., 2020 or **(Right panel)** two years post-surgery in the study reported by Petrus et al., 2018 (E) *PCOLCE* mRNA levels from human patients designated metabolically healthy lean (MHL; n=19), metabolically healthy obese (MHO; n=18) and metabolically unhealthy obese (MUO; n=14). Data are presented as mean ± SEM. Statistical significance was determined by one-way ANOVA **(F)** scRNA-seq of mouse WAT showing *Pcolce* expression among various cell types **(G) (Left Panel)** UMAP from public scRNA-seq dataset (Nahmgoong, Hahn, et al., 2022) showing mouse adipose stem cells from epididymal adipose tissue (EAT) and inguinal adipose tissue (IAT) from three dietary conditions: normal standard diet (SD), 2-week Western diet (2wWD), and 10-week Western diet (10wWD). Colors indicate tissue origin and dietary condition. **(Right panel)** UMAP of *Pcolce2* gene expression with dietary conditions. **Note:** *PCOLCE /Pcolce* scRNA-seq data of human **(A)** and mouse **(G)** were obtained from the single cell portal (Broad Institute of MIT and Harvard; Tarhak et al., 2025), based on datasets from Emont et al., 2022. Adipogenesis data and UMAP of human scRNA-seq **(Band C)** were acquired from from the adipose tissue Knowledge portal (Zhong et al., 2025), which includes data from Massier et al., 2023; Hinte et al., 2024; and Reinisch et al., 2024.

**Supplementary Figure 2.**
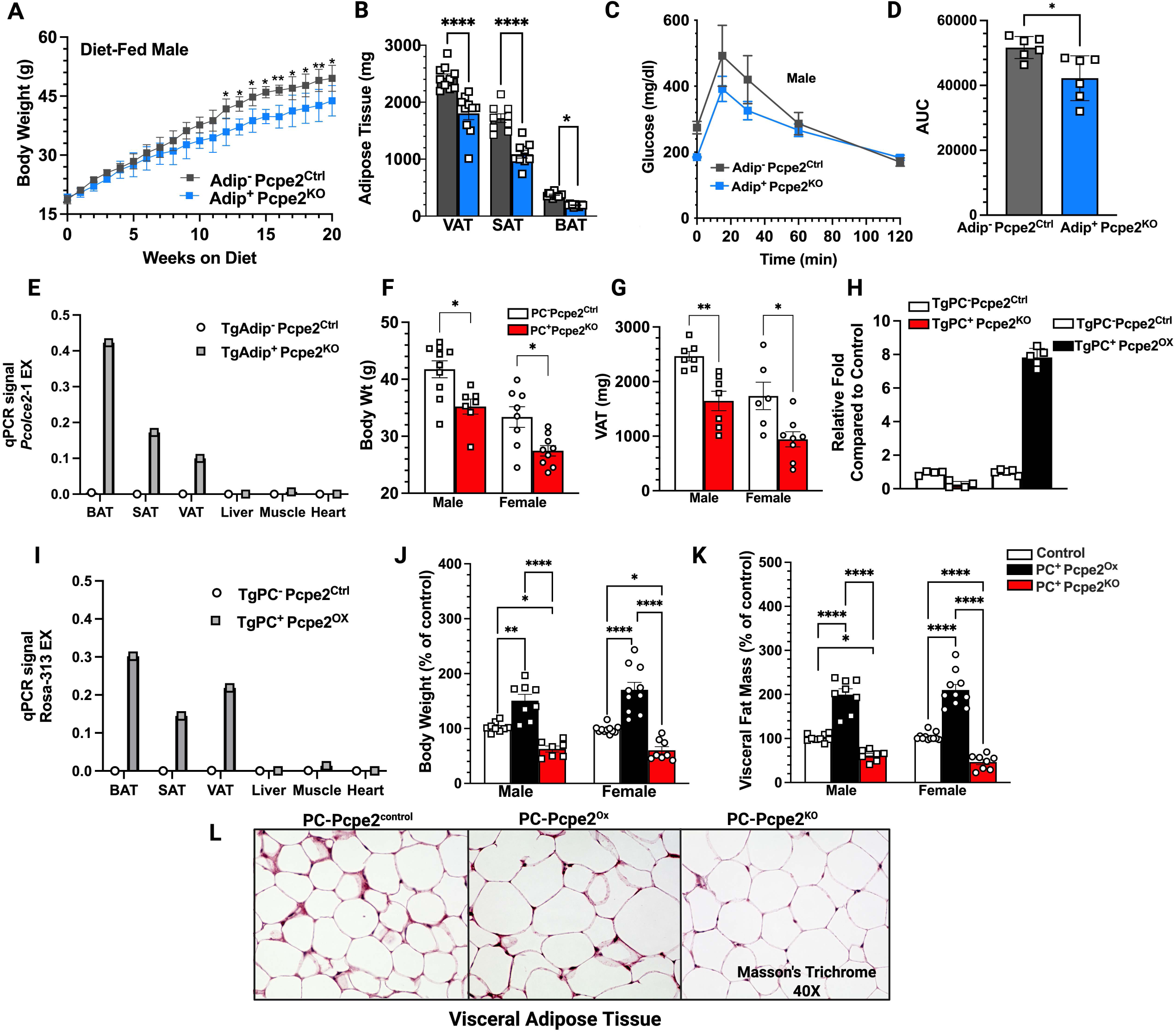
Pcpe2 Expression and Specificity of Cre Recombinase. Adip^+^Pcpe2^KO^ and Adip^-^Pcpe2^Ctrl^, control mice were fed a WO for 20 weeks. **(A)** Body weight of WO male mice from indicated genotype **(B)** End point body weight in male mice **(C)** Glucose tolerance test (GTT) in WO male mice as compared to their controls. **(D)** Area under the curve (AUC) for GT **(E)** Indicated mouse tissues were submitted to Transnetyx Genotyping Services for DNA analysis using specific probes developed for each mouse strain. The qPCR signal for the probe detecting *Pcolce2* floxed exon 3, *Pcolce2-1* EX, targets gene sequence after Cre recombinase removes exon 3 of the *Pcolce2* gene, thus, a positive signal indicates removal of the region indicated the specificity of the adiponectin Cre recombinase. Similar tissue specificity were noted when *Pcolce2* floxed mice were crossed with the Pdgfra (PC) Cre (data not shown) **(F-G)** Body and VAT weights in 20- weeks WO-fed male and female mice comparing TgPC+Pcpe2^KO^ and their negative littermate controls. Showing similar weight changes regardless of Cre recombinase **(H)** VAT Pcpe2 mRNA abundance from male mice fed SD from indicated genotype, shows Pcpe2 overexpression in TgPC+Pcpe2^Ox^ mouse stain compared to its control (I) Indicated mouse tissues were submitted to Transnetyx Genotyping Services for DNA analysis using specific probes developed for each mouse strain. Transgenic mice expressing Pdgfra (PC+) were crossed with *Pcolce2* overexpressor mice, TgPC +Pcpe2^Ox^ mice, and their negative PC^-^ controls PC^-^Pcpe2^Ctrl^ For this strain the PCR signal is plotted for the probe detecting Rosa-313 EX, which targets the floxed DNA sequence corresponding to the stop codon which is removed after Cre recombinase allowing overexpression of the *Pcolce2* gene. Thus, a positive signal indicates that the stop codon has been removed **(J-K)** Body and VAT weights expressed as percent of control (PC^-^ littermate control used for each mouse strain) from 20-25 week WO fed male and female mice of the indicated genotype. **(L)** Representative images of VAT sections stained with Masson’s Trichrome from indicated genotype. All data represent the mean ± SEM (n=5-15). Statistical significance were analyzed using unpaired t-test or one-way ANOVA followed by Tukey’s post hoc test. *p :5 0.05.

**Supplemental Figure 3.**
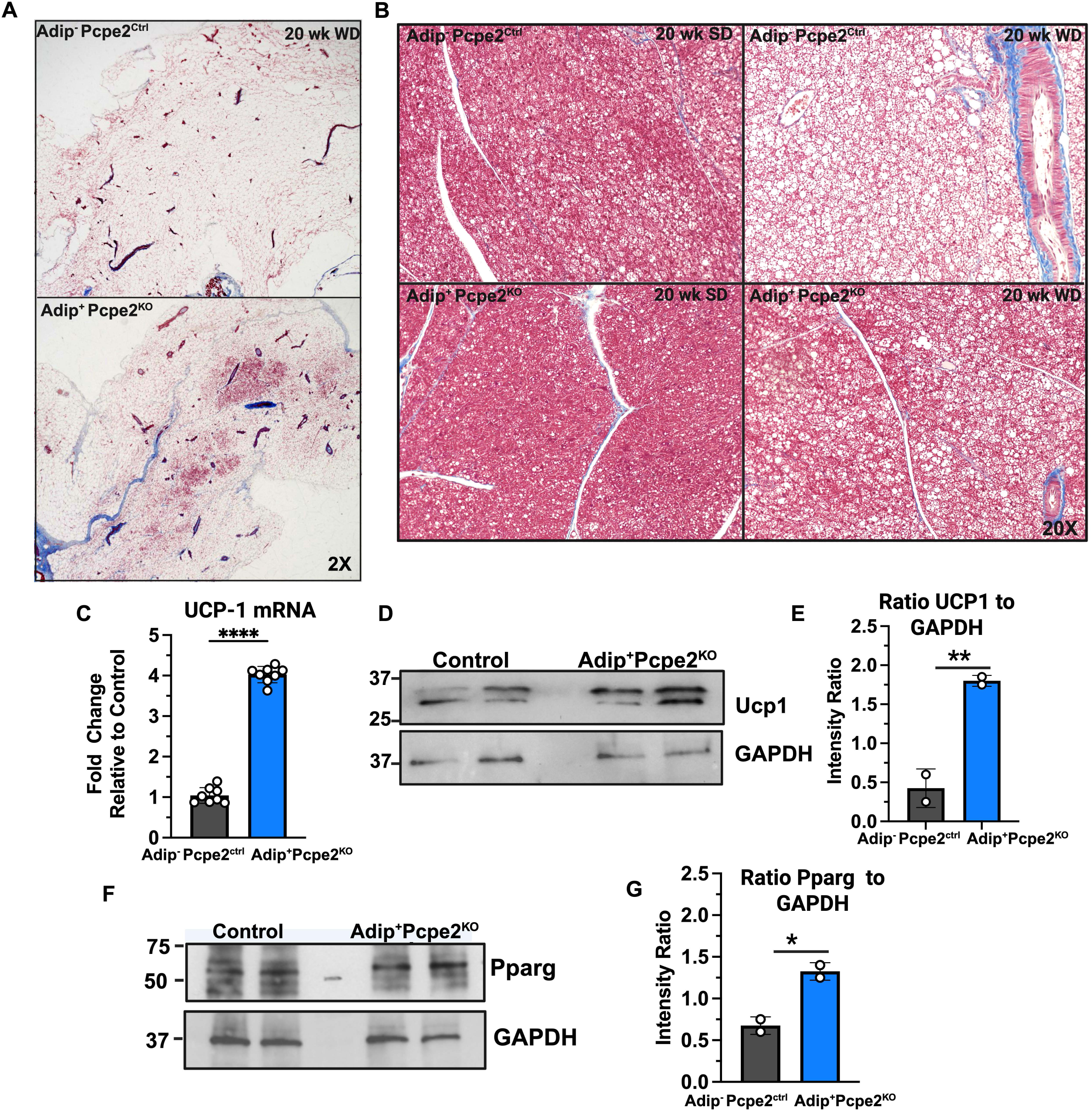
Absence of Pcpe2 Promotes WAT Beiging and Attenuates BAT Whitening. **(A)** Representative images of Masson’s Trichrome stained WAT 2X images from Adip·Pcpe2^Ctrl^**(top panel)** and Adip+Pcpe2^KO^ **(bottom panel)** mice fed WD for 20 weeks **(B)** Representative Masson’s Trichrome stained BAT images from Adip· Pcpe2^Ctrl^ **(top panels)** and Adip+Pcpe2^KO^ **(bottom panels)** mice fed SD **(left panels)** or WD **(right panels)** for 20 weeks **(C)** UCP1 mRNA abundance in SAT of indicated mouse genotype **(D)** Western blot analysis of Ucp1 and glyceraldehyde phosphate dehydrogenase (GAPDH) from SAT of WO-fed mice of indicated genotype **(E)** Quantification of band intensity as a rato of UCP1 to GAPDH **(F)** Western blot analysis of Pparg and GAPDH from SAT of WO-fed mice of indicated genotype **(G)** Quantification of band intensity as a ratio of Pparg to GAPDH. PCR data represent mean± SEM (n=8) Statistical significance was determined by unpaired t-test. Western blot lanes show protein extracts from two different animals of each genotype.

**Supplemental Figure 4.**
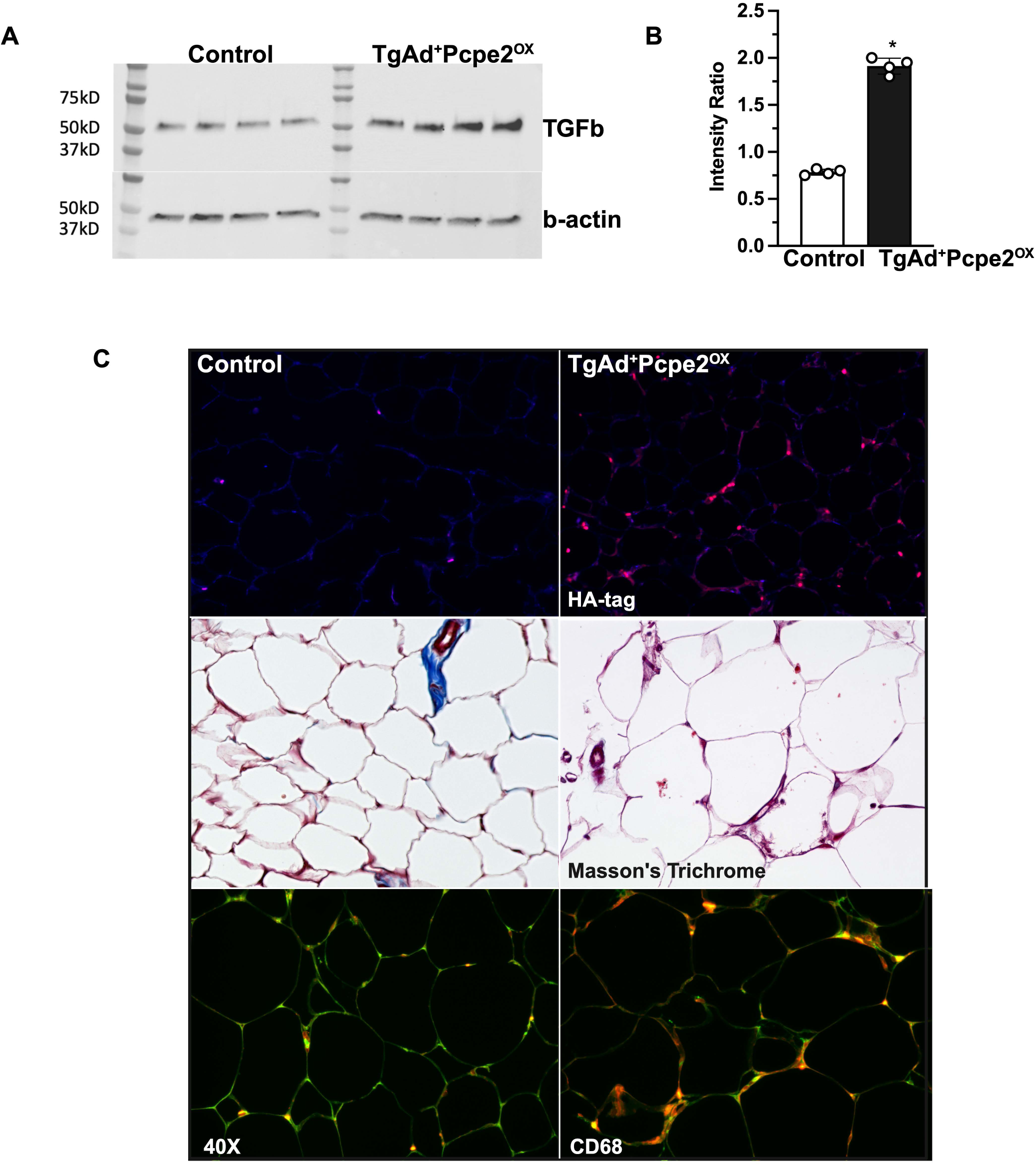
Pcpe2 Overexpression Enhances TGFb Signaling and Inflammation in VAT. FAPs were isolated from VAT SVF using collagenase digestion and then isolated by FAGS sorting and cultured for 4 days. Cell were treated with 2 ng/ml of TGFb1 for 90 min after which the culture medium was collected and subjected to Western analysis with intensity ratio quantification **(A)** Sections of re-harvested transplanted VAT were stained for hemagglutinin antigen (HA) to detect the fusion protein Pcpe2-HA tag expressed in TgAd+Pcpe2^Ox^ mice **(top panels),** Masson’s trichrome **(middle panels),** CD68 and perilipin **(bottom panels).**

**Supplemental Figure 5.**
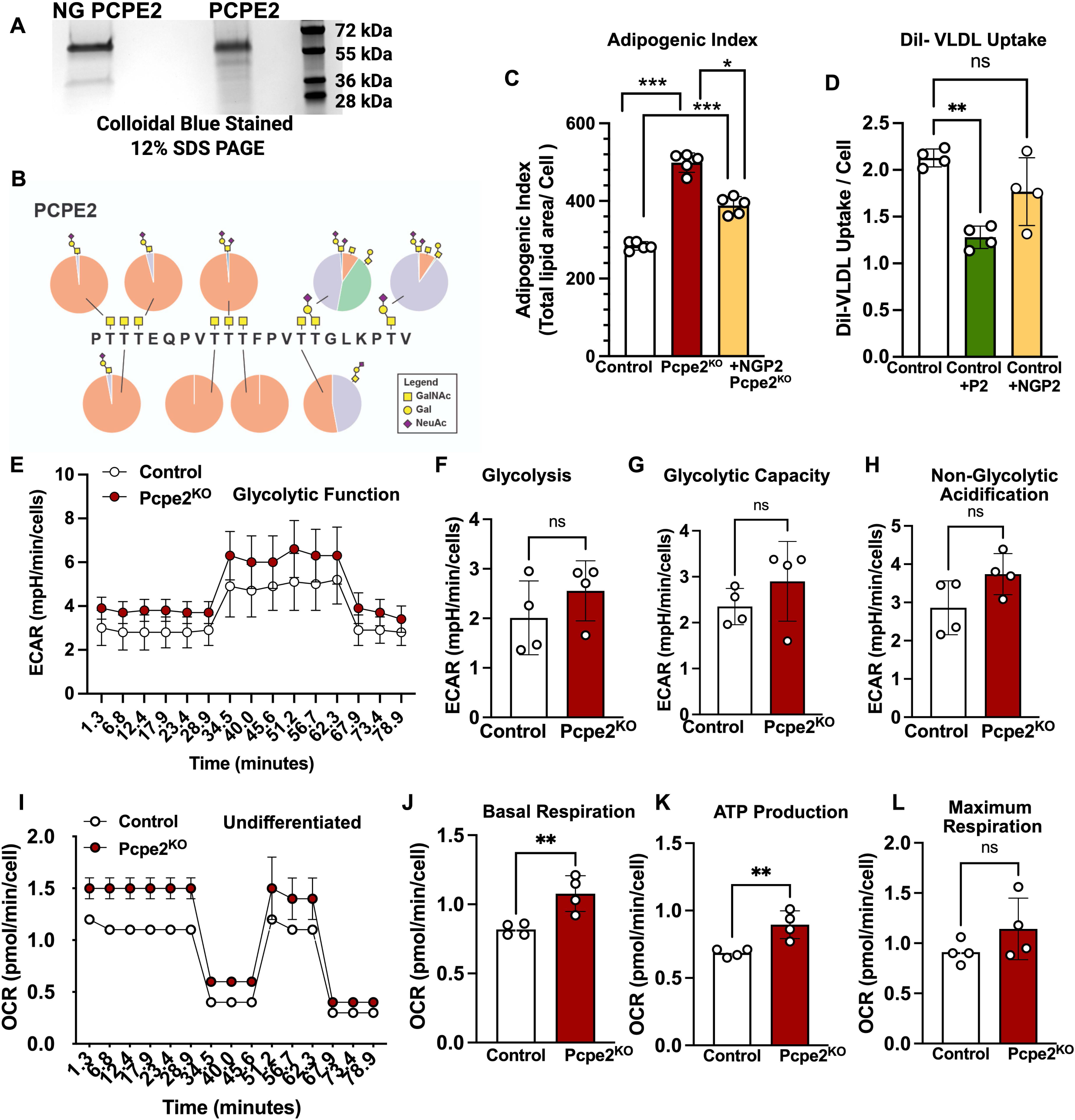
PCPE2 Glycosylation and Adipogenic Index. **(A)** PCPE2 and NG PCPE2 were purified from ExpiCHO expression system and aliquots applied to 12% SOS PAGE, then stained with colloidal blue **(B)** Mass spectrometry analysis showing glycans associated with indicated amino acids within linker region. Pie chart shows relative abundance for each 0-glycan detected **(C)** Adipogenic index from *ex vivo* differentiated adipocytes from control or Pcpe2^KO^ mouse SAT SVF **(D)** Dil-VLDL uptake in differentiated adipocytes from indicated genotype normalized to total cell number per well **(E)** Seahorse study showing extracellular acidification rate traces from differentiated adipocytes from control and Pcpe2^KO^ SAT SVF **(F)** Glycolysis **(G)** Glycolytic capacity **(H)** Non-glycolytic acidification (I) Seahorse study showing oxygen consumptKon rate curve from undifferentiated SVF cells derived from control and Pcpe2^KO^ SAT SVF **(J)** Basal Respiration **(K)** ATP Production **(L)** Maximum Respiration. All data represent mean ± SEM of differentiated adipocytes from SAT SVF with n=6 wells per genotype -96 well plate. Statistical significance was determined by unpaired t-test or ANOVA.

## Notes

### Competing Interest Statement

The authors have declared no competing interest.

